# Post-outbreak surveillance reveals contrasting circulation dynamics of West Nile, Usutu and Bagaza viruses in southern Portugal, 2021–2023

**DOI:** 10.64898/2026.09.21.752589

**Authors:** Catarina Fontoura-Gonçalves, David Gonçalves, Luís P. da Silva, Francisco Llorente, Alberto Moraga-Fernández, Marinela Contreras, Tatiana Silva, Tereza Almeida, Catarina J. Pinho, Sara Minayo-Martín, Victor Rodriguez-Valencia, Manuella García-Fasanello, Ana Margarida Lopes, Joana Abrantes, Gonçalo de Mello, João Basso Costa, Paulo Célio Alves, Elisa Perez-Ramirez, David Roiz, Miguel Ángel Jiménez-Clavero, Ursula Höfle, João Queirós

**Author notes:** **Correspondence:** Catarina Fontoura-Gonçalves and João Queirós.

## Abstract

Mosquito-borne orthoflaviviruses pose a significant threat to human and animal health worldwide. In Europe, West Nile (WNV), Usutu (USUV) and Bagaza (BAGV) viruses have expanded their geographic distribution and are detected with increasing frequency. Following the emergence of BAGV in 2021, we investigated the circulation dynamics and genomic diversity of orthoflaviviruses in southern Portugal, where their epidemiology remains poorly understood. Active *Orthoflavivirus* surveillance was conducted in birds and mosquitoes during 2021-2023 using serological and molecular assays, complemented by viral genome sequencing and phylogenomic analyses. Viral RNA was detected in 3.5% (41/1,171) of birds, while seroneutralization assays identified an overall seroprevalence of 55.3% (503/910). WNV RNA (lineage-1, strains WMed-1.2 and WMed-1.3) was detected annually in resident birds, including *Alectoris rufa*, *Cyanopica cooki* and *Garrulus glandarius* (1.5%, 18/1,171), and in *Culex univittatus* mosquitoes (0.8%, 2/255). Together with a WNV-specific seroprevalence of 28.5% (259/910) in birds, these findings support sustained local circulation. USUV RNA (Africa 3.1 sub-lineage) was detected in *A. rufa* in 2021 and 2023 (0.2%, 2/1,171), but seroprevalence was low (1.3%, 8/623), consistent with intermittent circulation. BAGV RNA (genotype-2) was detected only during the 2021 outbreak in *A. rufa* and *Emberiza calandra* (1.8%, 21/1,171), although BAGV-specific antibodies (6.6%, 60/910) were detected for up to two years after the outbreak, suggesting continued exposure or low-level circulation. Overall, our findings reveal contrasting circulation dynamics of orthoflaviviruses in southern Portugal. Sustained WNV circulation implies the need to strengthen avian and human surveillance in Portugal to better assess its public health risk, while the sporadic detection of USUV and BAGV underscores the value of long-term, integrated One Health surveillance for improving early warning and preparedness for emerging mosquito-borne diseases.

## 1. Introduction

West Nile (*Orthoflavivirus nilense*, WNV), Usutu (*Orthoflavivirus usutuense*, USUV) and Bagaza (*Orthoflavivirus bagazaense*, BAGV) viruses are emerging mosquito-borne orthoflaviviruses of increasing relevance to human and animal health (Worden-Sapper et al., 2025). These RNA viruses belong to the genus *Orthoflavivirus,* family Flaviviridae, and are maintained in nature primarily through an enzootic transmission cycle involving mosquitoes as vectors and birds as the main reservoir hosts. Incidental infection can occur in other vertebrate hosts, including humans, although evidence of BAGV infection in humans remains limited and such hosts generally act as dead-end hosts (Benzarti et al., 2019). Over the last decade, increased epidemic activity, accompanied by a rise in human morbidity and mortality, has made mosquito-borne orthoflaviviruses an increasing public health concern across Europe (Rodríguez-Alarcón et al., 2021; Caro-Leiro et al., 2026). However, their circulation dynamics remain poorly characterized in Portugal, a country located at the southwestern edge of mainland Europe and the Iberian Peninsula, in close proximity to Africa and along major migratory bird routes connecting the two continents.

WNV belongs to the Japanese Encephalitis (JE) virus serocomplex, with lineage-1 and lineage-2 circulating in Europe (Koch et al., 2024). Within the Iberian Peninsula, knowledge of WNV circulation differs markedly between Spain and Portugal. In Spain, WNV has been detected in a wide range of hosts and vectors, including competent *Culex* spp. (Benzarti et al., 2019; Caro-Leiro et al., 2026), and major human outbreaks occurred in 2020 and 2024 (Rodríguez-Alarcón et al., 2021; Caro-Leiro et al., 2026). In neighbouring Portugal, however, the public health impact and circulation dynamics of WNV remains poorly understood, despite serological evidence of circulation in wild birds (Formosinho et al., 2006; Barros et al., 2011; Fontoura-Gonçalves et al., 2025), horses (Barros et al., 2017) and humans (Rocha et al., 2025). Molecular evidence of WNV circulation also remains scarce, with few genomic characterizations so far (WNV lineage-1): three in mosquitoes, in 1971 (Filipe, 1972), 2004 (Esteves et al., 2005) and 2025 (Gutiérrez-López et al., 2026), and one in a wild *Astur gentilis* in 2024 (Maroco et al., 2025).

The emergence of USUV in Europe dates to 1996 in Italy, a large epizootic in 2001 in Austria and subsequent spread to central Europe, with recurrent outbreaks in wild birds (Vilibic-Cavlek et al., 2020). USUV is mainly transmitted by *Culex* sp. but has been detected in a variety of mosquito species in Europe and Africa (Vilibic-Cavlek et al., 2020). In the last years, human cases have been progressively reported (Føh et al., 2026). In Portugal, the virus was first detected in 2021 (Africa 3.1 sub-lineage), in red-legged partridges (*Alectoris rufa*), and again in 2023 (Queirós et al., 2024). However, a seroepidemiologic study between 2018 and 2022 revealed low seroprevalence (2.1%, Fontoura-Gonçalves et al., 2025).

BAGV, synonymous to Israel Turkey Meningoencephalitis virus (ITV, Fernández-Pinero et al., 2014) belongs to the Ntaya serocomplex and was first detected in Central African Republic in 1966 in a pool of *Culex* spp. mosquitoes (Digoutte, 1978). In 2010, BAGV was detected for the first time in Europe, associated with an outbreak in *A. rufa* and *Phasianus colchicus* (Agüero et al., 2011). The virus re-emerged in Spain in 2019, causing a local outbreak in *A. rufa* (Höfle et al., 2022), followed by a larger outbreak in 2021 (Gonzálvez et al., 2024), which extended to Portugal, where BAGV was detected for the first time (Queiros et al., 2022). Despite primarily affecting *A. rufa*, the virus was also detected sporadically in other wild birds (Queiros et al., 2022; dos Santos et al., 2024; Sekee et al., 2024) and mosquitoes (*Anopheles artroparvus*, Sekee et al., 2024). Seroepidemiologic analyses revealed a sharp decline in BAGV circulation in the years following the outbreaks in *A. rufa* (García-Bocanegra et al., 2013; Fontoura-Gonçalves et al., 2025). Outside Europe, BAGV has been detected in domestic birds (Steyn et al., 2019) and in 17 mosquito species, belonging to different genera, with *Culex* spp. accounting for most detections (Sekee et al., 2024).

Wild birds play a pivotal role as sentinel species for orthoflaviviruses surveillance (Straková et al., 2015; Tamba et al., 2024), enabling early detection of viral circulation and potential spillover risk to humans. Among them, *A. rufa* has been shown to be highly susceptible to *Orthoflavivirus* infection and a competent host for WNV and BAGV (Pérez-Ramírez et al., 2018; Fontoura-Gonçalves et al., 2025). This galliform is widely distributed across Western Europe, ranging from introduced populations in the United Kingdom to native populations in France, Spain, Italy and Portugal, where it is of considerable ecological, economic and social importance. In this study, we investigated the epidemiological dynamics of orthoflaviviruses in *A. rufa* along with other bird species and mosquitoes after the 2021 Portuguese BAGV outbreak (Queiros et al., 2022).

## 2. Materials and methods

### 2.1 Study area, sampling, and data collection

The study was conducted between September 2021 and December 2023 in hunting estates in the municipalities of Mértola and Serpa in southern Portugal. Following the 2021 BAGV outbreak, a bi-monthly live-capture scheme targeting *A. rufa* was implemented in Serpa as an extension of the long-term integrated monitoring program established in 2018 (Fontoura-Gonçalves et al., 2025). Wild *A. rufa* were captured using 30 cage traps baited with wheat. Traps were activated during the morning and afternoon and equipped with signalling devices to indicate trap closure, allowing birds to be retrieved promptly and minimizing the time spent in the traps. In addition to live birds, *A. rufa* specimens found dead in the field or harvested during the hunting season (October-February) were opportunistically sampled both in Serpa and Mértola.

Other bird species were also captured in Serpa and Mértola, primarily during spring and late summer (migration periods), to complement the targeted *A. rufa* surveillance. Birds were captured using baited walk-in traps and mist nets (Dunn & Ralph, 2004), which were checked regularly to minimize time in traps. All captured birds were identified to species level and ringed, and age and sex recorded whenever possible. Blood samples were left to clot and then centrifuged for serum separation and stored at –20°C. Growing feathers were also collected whenever possible. Growing feathers were plucked using tweezers and stored in RNA later at – 80°C. After sampling and ringing, all live-captured birds were released at the capture site. All procedures were conducted by experienced researchers following appropriate animal welfare practices (see ethical statement).

Mosquito surveillance started in August 2022. Sampling was performed every two months in Serpa and during spring and late summer in Mértola. Mosquitoes were captured using both UV/white light mini-CDC type traps (BG-Pro, Biogents, Germany) baited with carbon dioxide and home-made gravid traps constructed based on the Reiter-Cummings gravid trap (Fynmore et al., 2021). At each sampling event, five traps were placed near water sources and remained active between dusk and dawn for two consecutive nights. Collected mosquitoes were transported alive under controlled conditions (humidity/temperature) and stored at –80°C upon arrival at the laboratory until further processing.

### 2.2 Serological screening

Orthoflaviviruses antibodies screening was performed using a commercial competitive ELISA (Ingezim West Nile Compac, Gold Standard Diagnostics, Spain) designed for the detection of WNV antibodies, with cross-reaction against USUV and BAGV (Llorente et al., 2019). To differentiate WNV, USUV and BAGV specific antibodies, micro virus neutralization test (VNT) was used. The VNTs were performed in the BSL-3 laboratory at Centro de Investigación en Sanidad Animal (CISA-INIA, CSIC) according to Llorente et al. (2019). Seroepidemiologic data on *A. rufa* collected in 2021-2022 were previously published in Fontoura-Gonçalves et al. (2025). These previously reported data were incorporated into the present study to provide a continuous temporal framework, allow comparison with subsequent sampling periods and other bird species sampled, and integrate the serological findings with newly generated viral RNA detection data obtained by RT-qPCR. Remaining seroepidemiologic data for all subsequent sampling periods on *A. rufa* was obtained following the methodology previously described (Fontoura-Gonçalves et al., 2025). For the other bird species, due to the reduced sample volume that was possible to obtain in most individuals, we used a modified strategy. Positive and doubtful ELISA samples were screened using VNT against WNV, USUV and BAGV in parallel, using serial twofold serum dilutions, routinely from 1:10 to 1:1280, with further dilutions performed when required to distinguish between viruses (up to 1:10240 in this study). Negative ELISA samples were analysed by VNT against BAGV at dilutions 1:10 and 1:20 and those samples with neutralizing antibodies were screened again by VNT against BAGV and USUV (ELISA negative samples were considered negative to WNV) in parallel using serial twofold serum dilutions (1:10-1:1280/10240). When VNT titres did not reach a four-fold difference between viruses, the result was considered inconclusive (Llorente et al., 2019) and the specificity for a given *Orthoflavivirus* could not be determined (“undetermined orthoflaviviruses”). The number of samples analysed for USUV was lower than for WNV and BAGV because insufficient serum volume remained to test other bird species at USUV VNT dilutions >1:20.

### 2.3 Morphological and molecular identification of mosquitoes

Mosquitoes were processed on freezing plates under a stereomicroscope and identified to species or genus level based on key morphological characteristics (Becker et al., 2010). Blood-fed females were processed individually for *Orthoflavivirus* detection, and the thorax and abdomen were separated to distinguish potential infection from virus detected in the blood meal. Males and unfed females were pooled in groups of up to 20 individuals according to sex, genus/species, capture date, and sampling region.

To confirm the morphological identification of blood-fed females, we amplified the 658 bp Cytochrome c oxidase subunit I (COI) gene fragment using the primers LCO1490 and HCO2198 designed by Folmer et al. (1994) and following an adapted PCR protocol from the one described in Folmer et al. (1994): initial denaturation at 95 °C for 15 minutes, followed by 7 cycles at 95 °C for 30 seconds, annealing starting at 53 °C for 30 seconds with a decrease of 0.5 °C per cycle, and 72 °C for 1 minute. Subsequently, 33 amplification cycles were carried out with denaturation at 95 °C for 30 seconds, annealing at 50 °C for 30 seconds, and extension at 72 °C for 1 minute. A final extension step was performed at 60 °C for 10 minutes. PCR products were visualized on a 2% agarose gel, and positive amplifications were purified and Sanger sequenced following Folmer et al. (1994). As this approach failed to amplify the 658bp fragment in 27 specimens, a next-generation sequencing approach was applied, combining two primer pairs: the BF3 (Elbrecht et al., 2019) and BR2 (Elbrecht & Leese, 2017) for an expected size of 418 bps and fwhF1 (Vamos et al., 2017) and a modified version of Ill_C_R (5’-GGNGGRTANACNGTTCANCC-3’; Shokralla et al., 2015) for an expected size of 325 bps. All primers contained Illumina overhang adaptors at the 5’ end. Amplification was performed using an initial denaturation at 95 °C for 15 minutes, followed by 35 cycles of denaturation at 95 °C for 30 seconds, annealing at 50 °C for the BF3-BR2 primer pair or 45 °C for the fwhF1-modified Ill_C_R primer pair for 90 seconds, and extension at 72 °C for 30 seconds, followed by a final step at 60 °C for 10 minutes. A second PCR was performed to incorporate the Illumina sequencing adapters using 5 µl KAPA HiFi HotStart ReadyMix, 1 µl of index mix, 2 µl of ultrapure water and 2 µl of the previous PCR products diluted 1:10. The cycling condition consisted of an initial denaturation at 95°C for 15 minutes, followed by 35 cycles at 95°C for 30 seconds, at 50°C for 60 seconds and 72°C for 45 seconds, with a final extension at 60°C for 10 minutes. These were cleaned using 0.8 × AMPure XP beads, quantified using Epoch, diluted to 15 nM and pooled by primer. The libraries were then quantified by qPCR and pooled to obtain a final library at 4 nM. The final library was sequenced on an Illumina MiSeq System, using a MiSeq V2 500-cycle reagent kit, with a coverage of approximately 5,000 paired-end reads per sample and primer pair.

The OBITools software was used to bioinformatically process the sequences (Boyer et al., 2016). Paired-end reads were first aligned using the illuminapairedend command, and those with an overlap quality of less than 40 were discarded. Unaligned sequences were also removed using obigprep command. Remaining sequences were dereplicated to unique sequences per sample using obiuniq. Primers were removed using ngsfilter, and sequence lengths were trimmed to the expected lengths mentioned above (418 and 325 bp). Sequences with fewer than 10 reads were also removed. Clustering analysis was performed using sumacluster, with a sequence similarity threshold of 97% (Boyer et al., 2016). The obtained sequences were analysed in Geneious Prime® 2024.0.3 by assembling the two amplicons and mapping them against the Portuguese mosquito DNA library database (Madeira et al., 2021), which was then further confirmed by BLASTn (Brister et al., 2015).

### 2.4 RNA extraction and orthoflaviviruses detection

Total RNA from growing feathers and mosquitoes was extracted using a MagMAX CORE Nucleic Acid Purification kit™ in the KingFisher Apex instrument™, including negative extraction controls. Samples were screened for orthoflaviviruses using a duplex quantitative reverse transcription PCR (RT-qPCR) for the simultaneous and differential detection of Japanese encephalitis (JE) and Ntaya serocomplex (Kogut et al., 2020). Positive samples to the JE serocomplex were further submitted to another RT-qPCR for the simultaneous detection of WNV lineage-1 and lineage-2 (Jiménez-Clavero et al., 2006). Positive JE samples, but WNV negative, were further screened for detection of USUV using a specific RT-qPCR for amplification of part of the non-structural protein 5 (NS5) gene region of USUV (Weissenböck et al., 2013). Ntaya serocomplex-positive samples were screened for BAGV using a conventional nested RT-PCR targeting part of the NS5 gene (Sánchez-Seco et al., 2005) and following the procedures previously described (Queirós et al., 2022).

### 2.5 Whole genome sequencing and phylogenomic analysis

A subset of RT-qPCR-positive samples to WNV and BAGV were selected for whole-RNA library preparation based on virus, bird species, sampling year, and sampling month, aiming to capture the diversity of viral detections across host species and sampling periods. RNA concentration and quality were assessed with a Qubit 4™ Fluorometer (RNA HS Assay Kit™) for precise quantification, and purity RNA concentration was confirmed via NanoDrop™ Lite Spectrophotometer (Thermo Fisher Scientific, MA, USA), examining A260/A280 and A260/A230 ratios. Residual genomic DNA was removed by DNase I treatment (NZYtech, Portugal), and RNA was further purified using the GeneJET RNA Purification Kit (Thermo Scientific, MA, USA). RNA integrity and concentration were re-evaluated on an Agilent 4200 TapeStation, with RIN values measured following manufacturer’s guidelines. Ribosomal RNA was removed using the NEBNext rRNA Depletion Kit v2 (Human/Mouse/Rat; New England Biolabs), and the rRNA-depleted RNA was used to prepare libraries with the NEBNext ® Ultra TM II RNA Library Prep Kit (New England Biolabs) following the manufacturer’s protocol. Sequencing was conducted on a NovaSeq 6000 platform (150 bp paired-end) at Macrogen (Seoul, Korea). The genome consensus sequences were constructed using Geneious Prime® 2024.0.3. The paired-end data were combined for each sample and subsequently cleaned using BBDuk pluggin and merged using BBmerge. The sequences were then mapped against their reference genomes (NC_009942.1 for WNV; NC_012534.1 for BAGV, retrieved from NCBI) using default features with medium/low sensitivity 5x and do not trim function and deposited in the GenBank database.

Phylogenomic analyses were performed using the genomes generated in this study for WNV and BAGV, together with publicly available sequences retrieved from the NCBI Virus database (https://www.ncbi.nlm.nih.gov/labs/virus/). For WNV, publicly available sequences under the terms “West Nile virus” were retrieved on 17 February 2025 (n=76) and downloaded into Geneious Prime® 2024.0.3. Representative sequences of WNV lineages 1a (n=56) were selected for further analyses. For BAGV, sequences under the terms “Bagaza virus” and “Israel turkey meningoencephalitis virus” (ITV) were retrieved on the same date. ITV was included in the analyses as BAGV and ITV are genetically close and recognized by some authors as the same viral species (Fernández-Pinero et al., 2014). BAGV complete sequences (n= 34) were aligned in MAFFT and cropped to the coding region (10,281 bp) of the BAGV reference genome (NC_012534.1). Cropped sequences were realigned with complete retrieved ITV sequences, resulting in a total of 35 (29 BAGV and 6 ITV) sequences. ModelFinder from IQTree was used to select the best-fitting models according to BIC (Kalyaanamoorthy et al., 2017). Maximum likelihood phylogenomic trees were constructed with IQ-TREE web server (Trifinopoulos et al., 2016) using TIM+F+G4 and 1,000 replicates of ultrafast bootstrap resampling (Hoang et al., 2018) and SH-aLRT test (Minh et al., 2020). The consensus tree was visualized in Itol (Letunic & Bork, 2024) and further edited in Inkscape v1.3.2. Additionally, pairwise nucleotide identity was calculated for the partial genomes over the overlapping aligned regions with complete genomes using Geneious software (Geneious Prime® 2024.0.3). USUV sequences were not re-analysed here, as their phylogenomic characterization was previously reported (Queirós et al., 2024).

## 3. Results

### 3.1 Birds and mosquitoes sampled

In total, 1,367 birds (Figure 1) from 52 species were captured, from which 1,171 growing feathers and 1,070 blood samples were collected (Suppl. Table S1 and S2). *A. rufa* was the most represented species, with 893 samples, mostly from the bi-monthly monitoring program in Serpa (n=813, Suppl. Table S3). Among the other captured birds, the most representative families were Columbidae (n=140), Fringillidae (n=88) and Corvidae (n=57; Figure 1).

**Figure 1.**
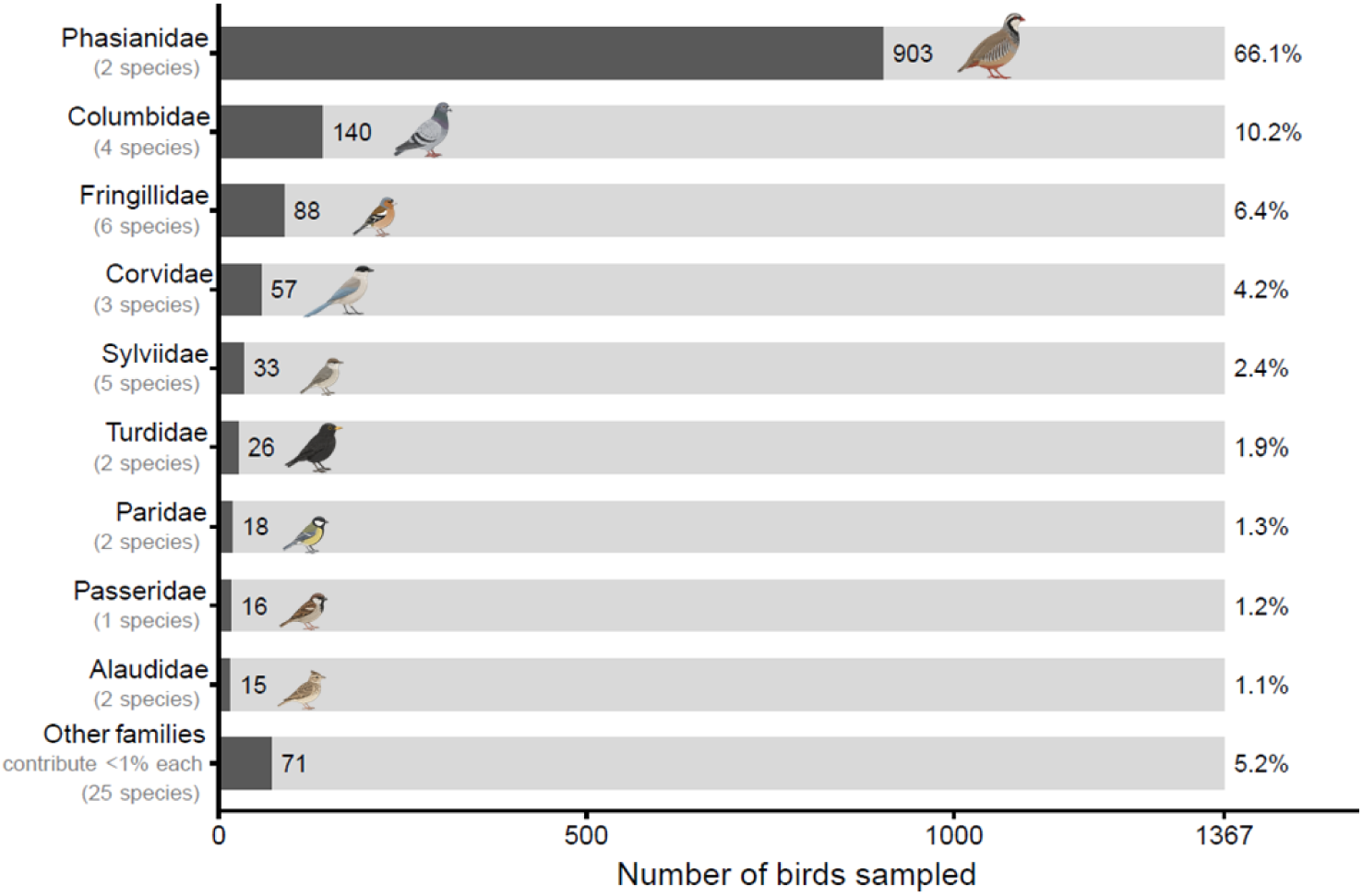
Number of individuals captured per bird family throughout the study period (2021-2023). The number of individuals is included after each dark bar, while the corresponding percentage (%) in relation to the total is indicated after each light grey bar. Bird illustrations were generated using the generative AI tool ChatGPT (OpenAI) and edited by the authors to depict representative species of each bird family.

Overall, 255 mosquitoes were collected and morphologically identified into 200 females (59 blood-fed and 141 unfed), 50 males and five of unknown sex. Females were taxonomically characterized into four genera and 15 species/complexes (Figure 2A, Suppl. Table S4). The predominant genus was *Culex* (87.0%), followed by *Aedes* (5.5%), *Anopheles* (5.0%) and *Culiseta* (2.5%). The most frequent species were *Cx. pipiens*/*torrentium* (17.0%) and *Cx. univitattus*/*perexiguus* (16.5%). Blood-fed females (n=59) were mainly captured in June 2023 (24.0%) and October 2023 (42.0%), and most (95.0%) were molecularly identified (Figure 2B and Suppl. Table S5), confirming *Culex* sp. as the predominant genus (88.2%). The dominant species were *Cx. pipiens* (40.7%; with both subspecies present, *Cx. p. molestus* [3.4%] and *Cx. p. pipiens* [1.7%], the remaining *Cx. pipiens* specimens [35.6%] could only be identified to species level), *Cx. theileri* (17.0%), and *Cx. univittatus* (15.3%). The COI sequences obtained were deposited in GenBank database (Accession numbers: PZ338730-PZ338744; PZ364570-PZ364571; PZ383187-PZ383200; PZ385223-PZ385247).

**Figure 2.**
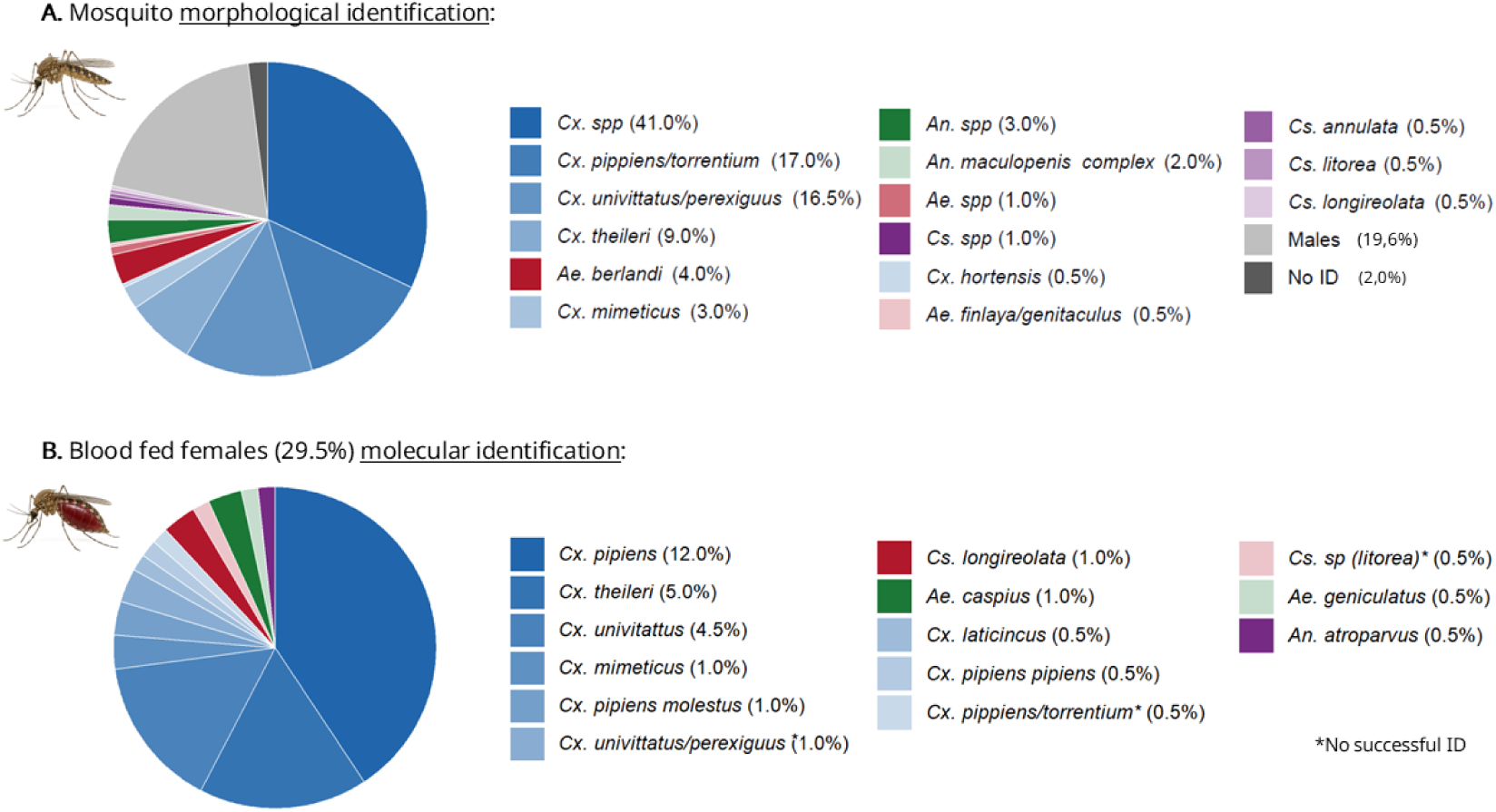
Mosquitoes collected and identified in this study. (A) Specimens morphologically identified by sex (n = 255) and taxa (only females, n = 200). (B) Blood-fed females (n = 59) were molecularly characterized using COI (Folmer et al., 1994). Mosquito illustrations were generated using the generative AI tool ChatGPT (OpenAI) and edited by the authors to depict representative mosquitos for each group.

### 3.2 ELISA and micro sero-neutralization test (VNT)

*Orthoflavivirus* seroprevalence was 46.4% (496/1,070) by ELISA and 55.3% (503/910) by VNT (Suppl. Table S2). Overall, VNT seroprevalence ranged from 85.0% (34/40) in November 2021 to 32.7% (33/101) in August 2022. For *A. rufa* in Serpa, seroprevalence was 71.8% (375/524), ranging between 94.6% (35/37) in September/October 2023 and 23.8% (10/42) in August 2022 (Suppl. Figure S1 and Table S3A).

WNV specific antibodies were found in 18 bird species, with an overall seroprevalence of 28.5% (259/910), ranging between 44.7% (38/85) in April 2023 and 21.7% (10/46) in December 2023 (Figure 3). For *A. rufa* in Serpa (n=524), seroprevalence ranged between 52.9% (18/34) in April 2023 and 9.5% (4/42) in August 2022, with a global seroprevalence of 31.9% (167/524; Suppl. Figure S2 and Table S3A). For the other most frequently captured species (>20 individuals), seroprevalence ranged from 83.3%, in *Coccothraustes coccothraustes,* to 16.1%, in *Turdus merula* (Suppl. Table S6). Seroprevalences by sampling periods for other non-Phasianidae species are present in Suppl. Table S7.

**Figure 3.**
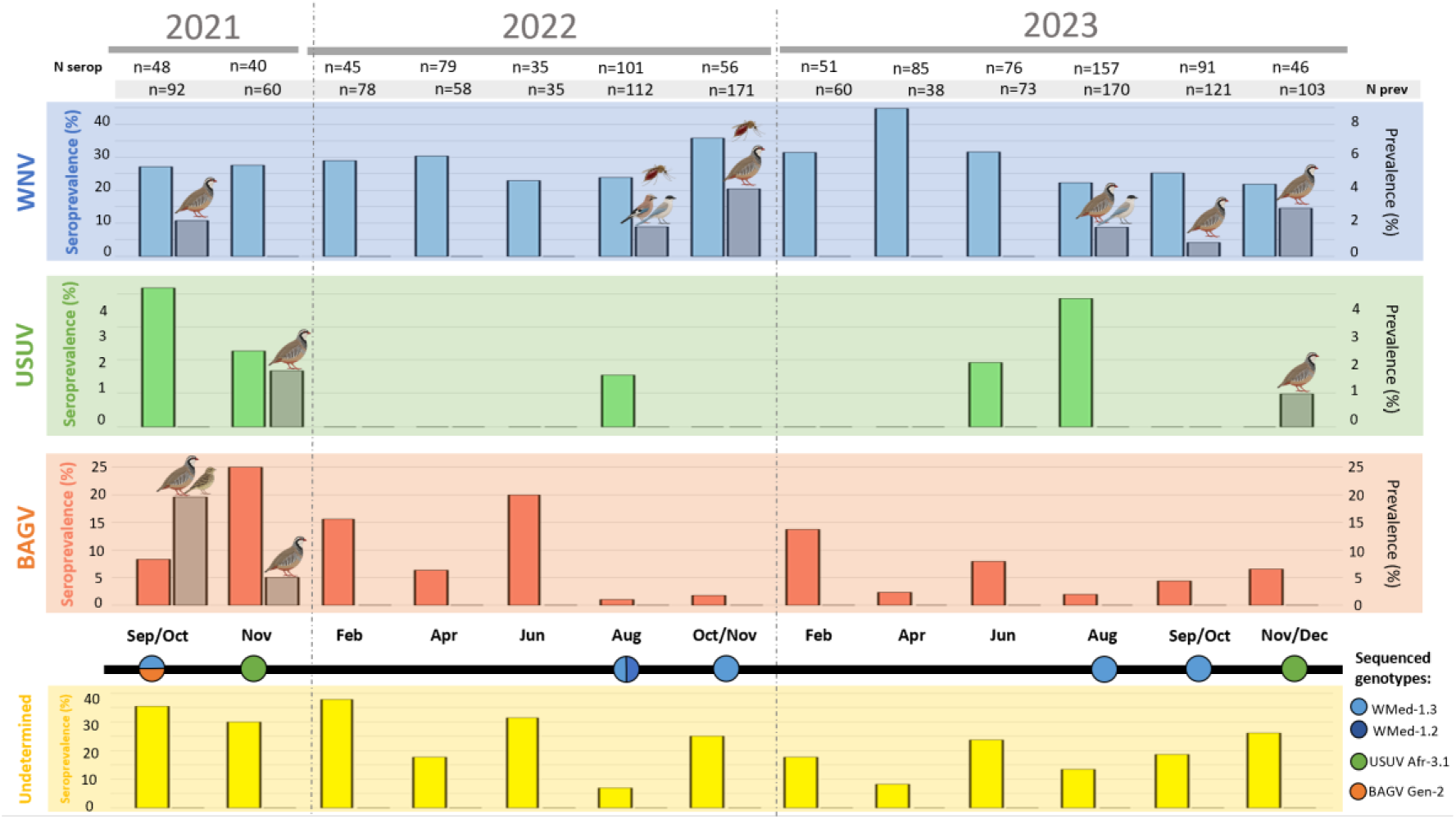
Temporal dynamics of West-Nile virus (WNV, blue), Usutu virus (USUV, green) and Bagaza virus (BAGV, orange), in avian hosts and mosquito vectors following the emergence of BAGV in Portugal, September 2021. Seroprevalence of virus-specific antibodies is shown on the left axis and represented by coloured bars, while the prevalence of virus detection is shown on the right axis and represented by grey bars. Seroprevalence of “undetermined orthoflaviviruses” antibodies is also displayed (yellow bars). The bird and mosquito species in which viral genome was detected are indicated on top of the bars: WNV was detected in *A. rufa*, *Garrulus glandarius*, *Cyanopica cooki* and *Culex univitattus*. USUV was detected in *A. rufa*. BAGV was detected in *A. rufa* and *Emberiza calandra*. The number of birds analysed is indicated above each sampling period (in black for seroprevalence and in grey for prevalence). In the chronological bold bar, colour-coded circles represent virus molecular detection, highlighting active co-circulation. Each colour corresponds to a specific genotype (light blue for WMed-1.3; dark blue for WMed-1.2; orange for BAGV genotype-2 and green for USUV African 3.1 sub-lineage). Bird and mosquito illustrations were generated using the generative AI tool ChatGPT (OpenAI) and edited by the authors.

USUV specific antibodies were found in four species, with an overall seroprevalence of 1.3% (8/623; Figure 3). In Serpa, only four *A. rufa* had specific antibodies (0.8%, 4/476; Suppl. Figure S3 and Table S3A). For the other bird species, USUV antibodies were found in three individuals, a *Parus major* in August 2022, a *Pica pica* in June 2023 and a *Lanius meridionalis* in August 2023 (Suppl. Table S7).

BAGV-specific antibodies were detected in *A. rufa* and one *Columba livia*, with an overall seroprevalence of 6.6% (60/910; Figure 3). For *A. rufa* in Serpa (n=524), these values ranged between 26.3% (10/38) in November 2021 and 2.3% (1/44) in October/November 2022, with a global seroprevalence of 11.3% (59/524; Suppl. Figure S4 and Table S3A).

Antibodies of undetermined specificity (“undetermined orthoflaviviruses”) were detected in 12 species, with a seroprevalence of 19.3% (176/910; Figure 3). For *A. rufa* in Serpa (n=524), these values ranged between 37.8 % (17/45) in February 2022 and 11.9% (5/42) in August 2022, with a global seroprevalence of 27.9% (146/524; Suppl. Figure S1 and Table S3A).

### 3.3 General and specific RT-qPCRs for Orthoflaviviruses detection

In birds, the overall *Orthoflavivirus* RNA prevalence was 3.5% (41/1,171) and ranged between 21.7% in September/October 2021 (20/92) and 0.8% in September/October 2023 (1/121; Suppl. Table S2). WNV RNA (lineage-1) was detected in *A. rufa* (15/883), *Cyanopica cooki* (2/36) and *G. glandarius* (1/6), between August and November, and across the three years of the study and regions, with an overall prevalence of 1.5% (18/1171; Figure 3). USUV RNA was detected in

*A. rufa* in November 2021 and 2023 (Figure 3), with an overall prevalence of 0.2% (2/1,171). BAGV RNA was detected in *A. rufa* (20/813) and *E. calandra* (1/3) in Serpa between September and November 2021 (Figure 3), with an overall prevalence of 1.8% (21/1,171). In *A. rufa*, these values ranged between 19.6% (18/92) in September/October 2021 and 3.5% (2/58) in November 2021 (Suppl. Table S3A).

In mosquitoes, the overall *Orthoflavivirus* RNA prevalence was 0.8% (2/255). WNV RNA (lineage-1) was detected in two blood-fed females of *Cx. univitattus* (2/33), one in the abdomen of a mosquito sampled in August 2022 (1/20), and the other in the abdomen and thorax of a mosquito sampled in October 2022 (1/25; Figure 3). BAGV and USUV were not detected in mosquitoes throughout the study period.

### 3.4 Genomes and phylogenomic analysis

Six complete WNV genomes, five from birds and one from mosquitoes and one BAGV genome from *A. rufa* were obtained. In addition, three partial WNV genomes, two from birds and one from mosquitoes, and one partial BAGV genome was obtained from *E. calandra* (Suppl. Table S8). Both complete and partial genomes were deposited in GenBank (Accession numbers: PZ377060-PZ377066; PZ392148-PZ392165; PZ392167-PZ392171; PZ400790-PZ400794; PZ400796-PZ400817; PQ672170-79). The complete and partial USUV genomes (Accession numbers: PQ677887; PQ672170–PQ672179) have been previously characterized as belonging to the African 3.1 sub-lineage (Queirós et al., 2024). In the present study, these detections were included in the overall analysis of orthoflaviviruses circulation and genomic diversity.

Among the five complete WNV genomes obtained from birds, four were from *A. rufa* (2021, 2022 and 2023) and one was from *C. cooki* (2023; Suppl. Table S8). Phylogenomic analysis clustered these strains within lineage-1a, cluster 2 and subgroup WMed-1.3 (Figure 4). The two incomplete genomes were obtained from *C. cooki* (2023, 7.5% coverage) and *G. glandarius* (2022, 7.1% coverage), with 99.3% and 97.6% pairwise nucleotide identity of overlapping regions with the WNV reference genome (NC_009942.1), respectively. The *C. cooki* sequences also showed 99.7% pairwise nucleotide identity with a Spanish strain (WMed-1.3 subgroup) obtained in 2016 (OM302315.1), whereas the *G. glandarius* sequences showed 99.9% pairwise nucleotide identity with a WNV strain (WMed-1.2 subgroup) obtained in Spain in 2022 from a *Cx. perexiguus* (OQ357820.1). In the case of mosquitoes, the complete WNV genome was obtained from the abdomen of the *Cx. univitattus* sampled in October 2022 (22A_Culex univitattus_Portugal_2022), while the partial genome (7563 bp, 68.6% coverage) was obtained from the abdomen of a *Cx. univitattus* sampled in August 2022. Phylogenomic analysis clustered the complete genome within subgroup WMed-1.3 (Figure 4), while pairwise distance analysis revealed a high degree of similarity (99.7%) between the complete and partial genomes.

**Figure 4.**
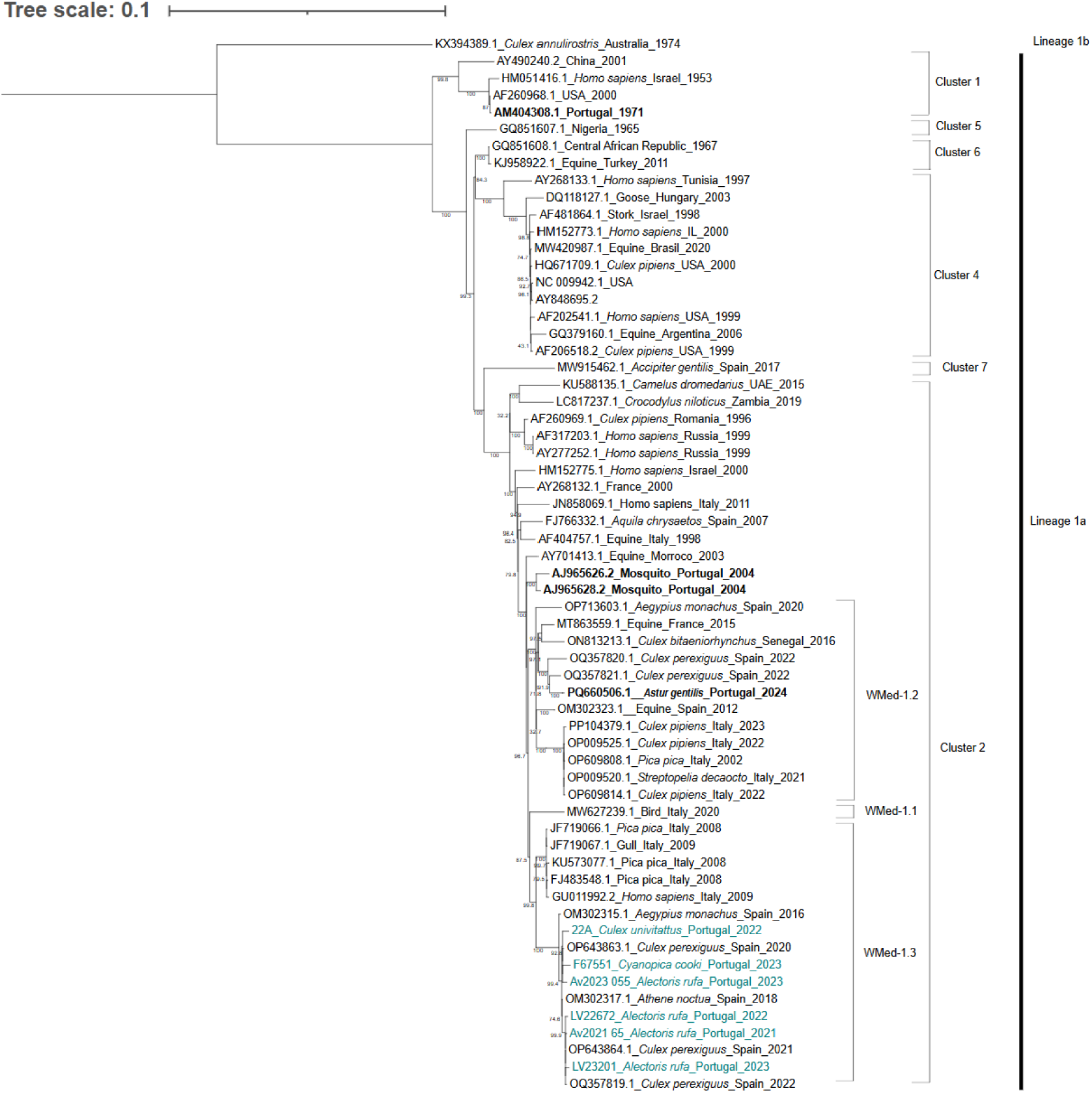
Maximum likelihood phylogenomic tree of the six complete protein-coding region genomes of WNV identified in this study (blue), combined with a subset of genomes representative of genetic diversity observed across strains within WNV lineage-1a, including those previously isolated in Portugal (bold). Tree was constructed with IQ-TREE web server using TIM+F+G4 and 1,000 replicates of ultrafast bootstrap resampling and SH-aLRT test. Tree scale is included on top. Bootstrap values are presented under the branches.

The complete (Av2021_008) and partial (Av2021_001) BAGV genomes were obtained from *A. rufa* and *E. calandra* found dead in October 2021, respectively. Pairwise distance analysis of overlapped regions (38.2% of the full genome) revealed a high degree of similarity between the two strains (99.8%). Phylogenomic analysis clustered the complete genome in BAGV-genotype 2 (Figure 5).

**Figure 5.**
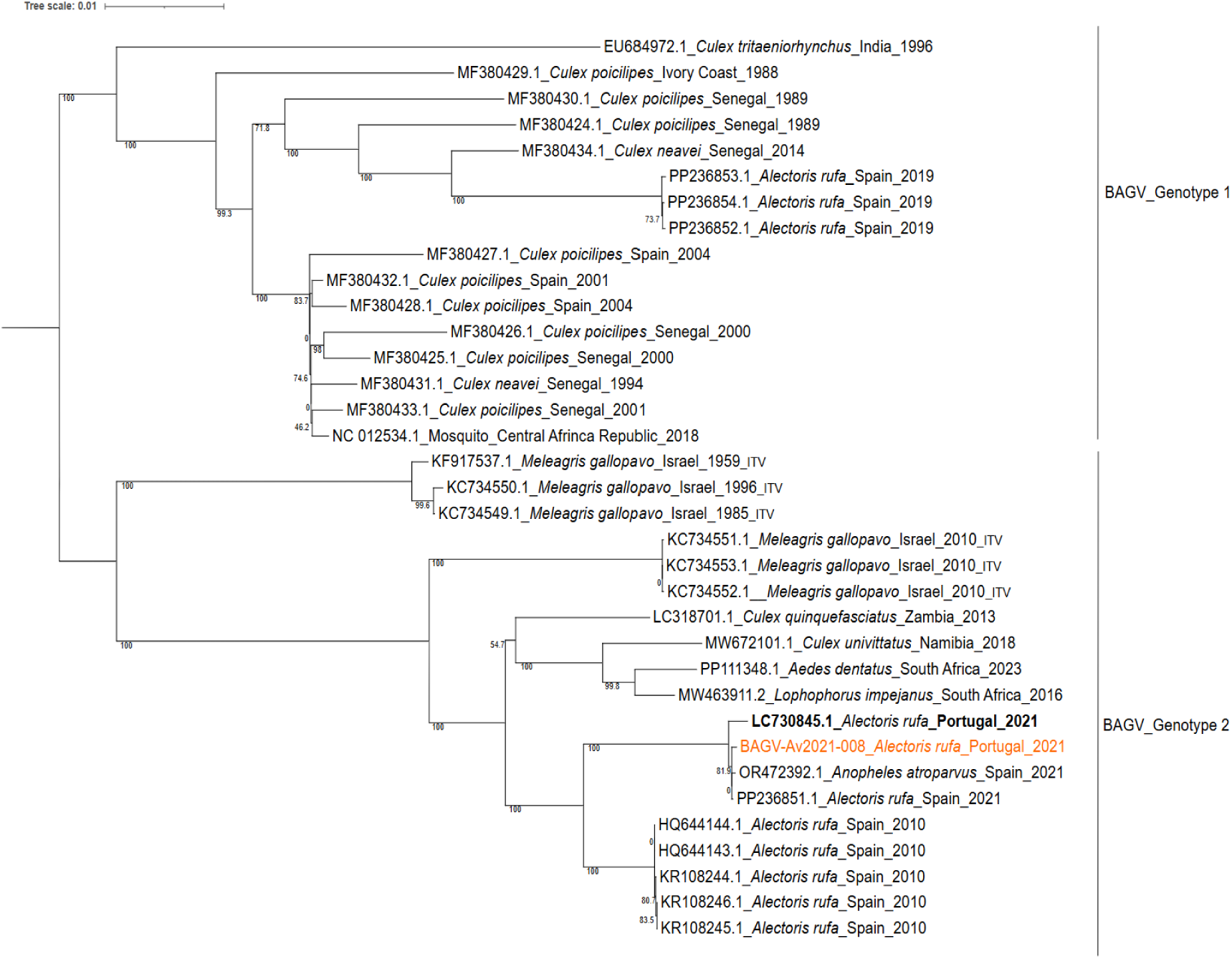
Maximum likelihood phylogenomic tree of the complete protein-coding region of the BAGV identified in *Alectoris rufa* found dead in October 2021 (orange), in comparison with the BAGV and Israel turkey meningoencephalitis virus (ITV) complete sequences available in NCBI (in bold BAGV previously described in Portugal). Tree was constructed with IQ-TREE web server using TIM+F+G4 and 1,000 replicates of ultrafast bootstrap resampling and SH-aLRT test. Tree scale is included on top. Bootstrap values are presented under the branches.

## 4. Discussion

By integrating viral surveillance, genomic analyses and longitudinal serological data from birds and mosquitoes, our study provides contrasting circulation dynamics of co-circulating WNV, USUV, and BAGV in mosquitos and avian hosts in southern Portugal. Located at the southwestern edge of Europe and in close proximity to Africa, this region is particularly exposed to the introduction and establishment of vector-borne pathogens, a process potentially intensified by climate change (Abbasi, 2026). Its position along a major Afro-Palearctic migratory bird flyways, together with the northward expansion of mosquitoes and their associated pathogens, further increases its epidemiological relevance in the European context (Aguilera-Sepúlveda et al., 2024b). Integrated epidemiological investigations are essential to identify drivers of pathogen emergence and characterize patterns of pathogen circulation in a One Health framework (Vilibic-Cavlek et al., 2020). When coupled with genomic analyses, these approaches can further elucidate pathogen origins, evolutionary histories, and transmission pathways (Maroco et al., 2025).

### Dynamics of zoonotic WNV and USUV

Our study provides new insights into the circulation dynamics of WNV and USUV in avian hosts and mosquito vectors in southern Portugal.

Albeit not related yet to human cases in the region, our findings indicate sustained WNV activity, supported by high annual seroprevalence in birds (21.7% – 44.7%) and repeated viral detections across multiple bird species and years. Together with the detection of WNV in mosquitoes, these findings provide strong evidence of endemic circulation in the region. WNV detections in birds and mosquitoes occurred between August and November, consistent with the seasonal pattern of orthoflaviviruses transmission described in southern Europe (Carrasco et al., 2024). Genomic analyses further revealed the co-circulation of WNV lineage-1 strains belonging to the WMed-1.2 and WMed-1.3 subgroups, consistent with the high diversity of WNV lineage-1 documented in Europe, including neighbouring Spain (Aguilera-Sepúlveda et al., 2024b). Strains belonging to the WMed-1.2 subgroup were first detected in Cádiz, Spain, in 2010 and rapidly spread throughout the country (Ruiz-López et al., 2023), with related strains also detected in Portugal (Maroco et al., 2025), supporting regional connectivity of WNV circulation across the Iberian Peninsula. WMed-1.3 strains were also detected in Spain in 2020 and were associated with one of the largest WNV outbreaks in humans reported in the country (Caro-Leiro et al., 2026). Importantly, strains belonging to this subgroup accounted for a substantial proportion of the WNV sequences detected in our study (Figure 4), raising questions about their epidemiological and potential public health relevance in Portugal. Moreover, the WMed-1.2 and WMed-1.3 strains identified here differ from those previously characterized in Portugal (Esteves et al., 2005; Filipe, 1972; Figure 4), suggesting temporal turnover and/or co-circulation of WNV strains and highlighting the importance of long-term surveillance integrating avian monitoring and viral genomics.

WNV RNA was detected in the abdomen and thorax of *Cx. univittatus* mosquitoes in 2022, providing further evidence of this species’ involvement in WNV circulation in the region. Indeed, *Cx. univittatus* is a recognized competent vector of WNV and was previously found WNV-positive in Portugal in 2004 (Esteves et al., 2005). Although primarily ornithophilic, this species has also been reported to feed on mammals, including humans, suggesting that it may act as a bridge vector facilitating enzootic transmission and spillover (Becker et al., 2010). In contrast, *Cx. perexiguus*, a species closely related to *Cx. univitattus,* has been proposed as an important vector in the WNV enzootic cycle in Southern Spain, with *Cx. pipiens* likely contributing to bridge transmission (Figuerola et al., 2022). These contrasting findings highlight the need to better characterize vector involvement and transmission dynamics in the region and to assess their implications for zoonotic spillover and human health in Portugal.

USUV RNA was detected only in *A. rufa* in November 2021 and November 2023, while seroprevalence remained low across bird species and years, supporting limited or intermittent circulation in southern Portugal. Both detections occurred in late autumn, which may indicate late-season USUV activity compared with other orthoflaviviruses that were generally detected between late summer and early autumn. However, this pattern should be interpreted cautiously given the limited number of USUV detections and may reflect differences in host competence, vector dynamics or sampling effort rather than distinct seasonality. Host competence may partly explain the low serological evidence of USUV circulation. Experimental infections indicate that *A. rufa* is a relatively inefficient amplification host for USUV compared with WNV (Llorente et al., 2023), consistent with the low USUV seroprevalence observed in our study. Nevertheless, repeated viral detection in *A. rufa* across two non-consecutive years suggests recurrent local USUV activity that may be difficult to detect through serological surveillance alone. Seroprevalence may also have been underestimated because serological screening was not optimized for USUV and may have missed low-titre antibodies, while VNT cross-reactivity with other orthoflaviviruses may have further hindered virus-specific antibody detection. USUV circulation in the region may therefore depend on other avian hosts, particularly *Turdus* and *Sturnus* species (Vilibic-Cavlek et al., 2020), which were insufficiently represented in our sample set. In this context, *A. rufa* may serve as a useful sentinel of local USUV activity without necessarily playing a major role in virus amplification, while other passerine species may contribute more substantially to viral maintenance. These findings underscore the importance of continued molecular surveillance across a broader range of avian hosts to better characterize USUV transmission dynamics in southern Portugal.

### BAGV outbreak investigation

Our results indicate that the BAGV outbreak detected in the *A. rufa* population in Serpa in 2021 was largely restricted to September – November. This timing resembles the first European BAGV outbreak reported in Spain in 2010, where cases were also detected during late summer and early autumn (García-Bocanegra et al., 2013). Despite continued molecular surveillance, BAGV RNA was not detected during the twelve sampling events performed between 2022 and 2023. Given our baseline sampling effort (26–40 *A. rufa* individuals per sampling period) and an assumed test sensitivity of 90%, we estimate a 95% probability of detecting at least one positive bird if true prevalence had been ≥8–12% (calculated using FreeCalc in EpiTools, Ausvet). During the peak transmission period (late summer and autumn), our sampling effort was substantially increased (42–159 individuals). This larger sample size significantly enhanced our detection power, lowering the detectable prevalence threshold to 8% (at the lower sampling effort of 42 birds) and 2% (at the maximum effort of 159 birds). The absence of BAGV detections during the periods after the outbreak therefore suggests that, if BAGV was circulating, prevalence remained below these detection thresholds. The mechanisms underlying such prolonged periods of low or undetectable virus activity, which may extend for nearly a decade, remain unclear, largely because of the scarcity of long-term longitudinal data on host exposure, vector abundance and virus circulation.

Phylogenomic evidence provides additional insight into possible epidemiological scenarios underlying BAGV emergence in Portugal. The BAGV strains isolated in Portugal were highly similar to genotype-2 strains detected in Spain in 2010 and 2021 (99.2% and 98.2%; Agüero et al., 2011). One possible explanation is the persistent, low-level circulation of BAGV genotype-2 within the Iberian Peninsula, followed by periodic amplification and geographic expansion. Alternatively, these findings are compatible with independent introductions of closely related genotype-2 lineages into Iberia from an unidentified African source (Aguilera-Sepúlveda et al., 2024a). The substantially lower sequence homology (92%) with the BAGV genotype-1 strain detected in Spain in 2019 (Aguilera-Sepúlveda et al., 2024a) further supports the possibility of recurrent introductions, potentially mediated by migratory birds and/or competent mosquito vectors, as previously proposed (Falcão et al., 2023; Aguilera-Sepúlveda et al., 2024a; Gonzálvez et al., 2024). These scenarios are not mutually exclusive: persistent local circulation and recurrent introductions may both contribute to BAGV epidemiology in Iberia. Long-term viral and genomic surveillance are required to disentangle their relative contributions.

Serological dynamics provide further evidence that BAGV exposure may have continued beyond the detectable 2021 outbreak. Seropositivity peaked approximately two months after the first viral detection (Fontoura-Gonçalves et al., 2025) and subsequently declined, with smaller increases observed in June 2022 and February 2023. Because the duration of antibody responses following BAGV exposure remains poorly characterized, these later increases could reflect either persistent antibodies following the 2021 outbreak or renewed virus transmission. Notably, one BAGV-seropositive partridge sampled in February 2023 (titre of 1:640) hatched in 2022, indicating exposure after April/May 2022 and therefore providing evidence of BAGV circulation after the initial outbreak. This interpretation is also consistent with a BAGV detection in a *P. pica* in Mértola in 2023 (dos Santos et al., 2024). Together, these observations suggest that the absence of BAGV RNA in our surveillance during 2022–2023 should not be interpreted as evidence of complete absence of virus circulation, but rather as compatible with continued activity at levels below our molecular detection capacity. Long-term capture-mark-recapture studies incorporating repeated serological sampling could help resolve antibody persistence at the individual level and further clarify BAGV transmission dynamics.

Although susceptibility to BAGV has been demonstrated in other bird species (Queiros et al., 2022; dos Santos et al., 2024), seroprevalence in non-galliform birds was markedly lower than in *A. rufa* (0.3%, 1/338). This difference may reflect lower exposure, reduced susceptibility, or weaker and/or less persistent antibody responses in other avian hosts. Given the apparently low level of virus circulation after the outbreak, detection in vectors may have been particularly challenging. Several factors could account for the absence of viral detections in mosquitoes, including low infection prevalence, limited sampling sensitivity, temporal mismatch between mosquito sampling and peak transmission, or involvement of vector species insufficiently represented in our sampling. A greater contribution of non-vector transmission routes may also be possible. Experimental studies have demonstrated direct bird-to-bird transmission of BGAV (Llorente et al., 2015), a route potentially amplified by high host densities and spatial aggregation around supplementary feeding sites, characteristic of intensively managed *A. rufa* hunting estates.

## 5. Conclusion

Our results revealed distinct eco-epidemiological patterns of WNV, USUV and BAGV in southern Portugal between 2021 and 2023. Sustained circulation of multiple WNV lineage-1 strains (WMed-1.2 and WMed-1.3) in birds, together with viral detection (WMed-1.3) in mosquitoes, provides strong evidence of endemic circulation and highlights the value of incorporating birds into active surveillance programs to better characterize transmission dynamics and provide early warning periods of increased spillover risk. The detection of WNV-positive *Cx. univittatus*, a recognized competent vector that may contribute to bridge transmission, further reinforces the need to better assess the public health implications of sustained WNV circulation and its potential impact on human and animal health. In contrast, USUV and BAGV showed more intermittent patterns of circulation and were detected molecularly only in wild birds. USUV was detected in two non-consecutive years but showed low seroprevalence, consistent with limited or intermittent local circulation. BAGV RNA detection was restricted to the 2021 outbreak in our surveillance; however, subsequent serological evidence, including antibodies in a bird hatched after the outbreak, together with later viral detection in the region (dos Santos et al., 2024) supports continued low-level circulation beyond 2021. These contrasting patterns illustrate how mosquito-borne orthoflaviviruses may exhibit markedly different transmission and evolutionary dynamics during co-circulation within the same ecosystem. Continued, integrated One Health surveillance combining longitudinal monitoring of birds and mosquitoes with viral genomics will be essential to resolve these transmission dynamics, strengthen early warning systems and improve preparedness for emerging orthoflaviviruses in Portugal, a strategically positioned region for the introduction and spread of mosquito-borne pathogens in Europe.

## Data availability

All data generated or analysed during this study are included in this preprint and the supplementary materials. Genomic sequence data have been deposited in the GeneBank database under accession numbers: PQ672170-79; PQ677887; PZ377060-PZ377066; PZ392148-PZ392165; PZ392167-PZ392171; PZ400790-PZ400794 and PZ400796-PZ400817 and are not yet made publicly available. Genomic sequence data will be released once the manuscript is published.

## Ethical statement

Legal permissions to capture and mark the animals were provided annually by *the Instituto da Conservação da Natureza e das Florestas* (ICNF; Portugal, Licences n° 49/2021, 54/2022, 53/2023, 510/2022/CAPT; 519/2023/CAPT, DGVF/DRCA/2021, DGVF/DRCA/2022 and DGVF/DRCA/2023). The study was evaluated by the Animal Welfare Ethics and Review Body at BIOPOLIS-CIBIO (reference no. 2024-10).

## Use of artificial intelligence tools

The authors acknowledge the use of ChatGPT (OpenAI) to generate the bird illustrations included in this manuscript. The authors reviewed, selected, and verified the final illustrations for scientific appropriateness.

## Acknowledgements

We thank all colleagues from Biopolis-CIBIO and IREC for their kind help with sample collection, storage, and processing. We are also grateful to the hunting states involved and their staff for their support and help in collecting the samples

