## Supplemental material for "Post-outbreak surveillance reveals contrasting circulation dynamics of West Nile, Usutu and Bagaza viruses in southern Portugal, 2021–2023"

**SUPPLEMENTARY MATERIAL**

This supplementary material is supporting information alongside the preprint “**Post-outbreak surveillance reveals contrasting circulation dynamics of West Nile, Usutu and Bagaza viruses in southern Portugal, 2021–2023**”.

**1. Supplementary Tables**

**Table S1.** List of birds captured by species and date. Numbers for the red-legged-partridge (*A. rufa*) are in bold.

| Year | 2021 |  |  | 2022 |  |  |  |  |  |  | 2023 |  |  |  |  |  |  |  | Total |  |
| --- | --- | --- | --- | --- | --- | --- | --- | --- | --- | --- | --- | --- | --- | --- | --- | --- | --- | --- | --- | --- |
| Season | Autumn |  |  | Winter |  | Spring | Summer |  | Autumn |  | Winter |  | Spring | Summer |  | Autumn |  |  |  |  |
| Month | Sep | Oct | Nov | Feb | Mar | Apr | Jun | Aug | Oct | Nov | Feb | Mar | Apr | Jun | Aug | Sep | Oct | Nov |  | Dec |
| *Actitis hypoleucos* |  |  |  |  |  |  |  |  |  |  |  |  |  |  | 1 |  |  |  |  | 1 |
| *Aegithalos caudatus* |  |  |  |  |  |  |  |  |  |  |  |  |  |  | 3 |  |  |  |  | 3 |
| *Alcedo atthis* |  |  |  |  |  |  |  |  |  |  |  |  |  |  | 6 |  |  |  |  | 6 |
| ***Alectoris rufa*** | **4** | **88** | **58** | **78** | **-** | **28** | **35** | **43** | **80** | **79** | **60** | **-** | **37** | **48** | **51** | **3** | **98** | **59** | **44** | **893** |
| *Anas platyrhynchos* |  |  |  |  |  |  |  |  |  |  |  |  |  |  |  | 2 |  |  |  | 2 |
| *Caprimulgus ruficollis* |  |  |  |  |  |  |  |  |  |  |  |  |  |  | 6 |  |  |  |  | 6 |
| *Linnaria cannabina* |  |  |  |  |  |  |  | 2 |  |  |  |  | 24 |  | 5 |  |  |  |  | 31 |
| *Carduelis carduelis* |  |  |  |  |  |  |  | 13 |  |  |  |  |  |  | 7 |  |  |  |  | 20 |
| *Chloris chloris* |  |  |  |  |  |  |  |  |  |  |  |  | 1 |  |  |  |  |  |  | 1 |
| *Certhia brachydactyla* |  |  |  |  |  |  |  |  |  |  |  |  |  |  | 1 |  |  |  |  | 1 |
| *Cettia cetti* |  |  |  |  |  |  |  |  |  |  |  |  |  |  | 1 |  |  |  |  | 1 |
| *Coccothraustes coccothraustes* |  |  |  |  |  |  |  | 3 |  |  |  |  | 22 |  | 2 |  |  |  |  | 27 |
| *Columba livia domestica* |  |  |  |  |  | 12 |  | 13 |  |  | 1 |  |  | 12 |  |  | 1 |  |  | 39 |
| *Columba palumbus* |  |  |  |  |  |  |  | 1 |  |  |  |  |  |  | 12 |  |  |  |  | 13 |
| *Cyanistes caeruleus* |  |  |  |  |  | 1 |  | 3 |  |  |  |  |  |  | 2 |  |  |  |  | 6 |
| *Cyanopica cooki* |  |  |  |  |  | 4 |  | 12 |  |  |  |  | 2 |  | 24 | 1 |  |  |  | 43 |
| *Dendrocopos major* |  |  |  |  |  |  |  | 1 |  |  |  |  | 1 |  | 6 |  |  |  |  | 8 |
| *Emberiza calandra* |  | 1 |  |  |  |  |  |  | 1 |  | 1 |  |  |  |  |  |  |  |  | 3 |
| *Ficedula hypoleuca* |  |  |  |  |  |  |  |  |  |  |  |  |  |  | 1 |  |  |  |  | 1 |
| *Fringilla coelebs* |  |  |  |  |  | 2 |  | 3 |  |  |  |  |  |  |  | 3 |  |  |  | 8 |
| *Galerida theklae* |  |  |  |  |  |  |  | 1 |  |  |  |  |  | 1 | 7 | 1 |  |  |  | 10 |
| *Gallus gallus domesticus* |  |  |  |  |  | 5 |  |  |  |  |  |  |  | 5 |  |  |  |  |  | 10 |
| *Garrulus glandarius* |  |  |  |  |  | 1 | 1 | 2 |  |  |  |  | 1 | 2 | 2 |  |  |  |  | 9 |
| *Hirundo rustica* |  |  |  |  |  |  |  |  |  |  |  |  | 1 |  |  |  |  |  |  | 1 |
| *Lanius meridionalis* |  |  |  |  |  |  |  | 1 |  |  |  |  |  |  | 2 |  |  |  |  | 3 |
| *Lanius senator* |  |  |  |  | 1 |  |  |  |  |  |  |  |  |  |  |  |  |  |  | 1 |
| *Lullula arborea* |  |  |  |  |  |  |  | 1 |  |  |  |  | 1 |  | 3 |  |  |  |  | 5 |
| *Luscinia megarhynchos* |  |  |  |  |  | 1 |  |  |  |  |  |  | 1 |  |  |  |  |  |  | 2 |
| *Merops apiaster* |  |  |  |  |  |  |  | 5 |  |  |  |  |  |  |  |  |  |  |  | 5 |
| *Oenanthe oenanthe* |  |  |  |  |  |  |  |  |  |  |  |  |  |  |  | 1 |  |  |  | 1 |
| *Oriolus oriolus* |  |  |  |  |  |  |  | 1 |  |  |  |  |  |  |  |  |  |  |  | 1 |

**Table S1** (continued). List of bird specimens captured by species and date. Numbers for the red-legged-partridge (*A. rufa*) are in bold.

| Year | 2021 |  |  | 2022 |  |  |  |  |  |  | 2023 |  |  |  |  |  |  |  | Total |  |
| --- | --- | --- | --- | --- | --- | --- | --- | --- | --- | --- | --- | --- | --- | --- | --- | --- | --- | --- | --- | --- |
| Season | Autumn |  |  | Winter |  | Winter | Winter |  | Winter | Winter |  | Spring | Summer |  | Autumn |  |  |  |  |  |
| Month | Sep | Oct | Nov | Feb | Mar | Apr | Jun | Aug | Oct | Nov | Feb | Mar | Apr | Jun | Aug | Sep | Oct | Nov |  | Dec |
| *Parus major* |  |  |  |  |  |  |  | 5 |  |  |  |  |  |  | 7 |  |  |  |  | 12 |
| *Passer domesticus* |  |  |  |  |  | 8 |  | 5 |  |  |  |  | 3 |  |  |  |  |  |  | 16 |
| *Phoenicurus phoenicurus* |  |  |  |  |  |  |  |  |  |  |  |  |  |  | 1 |  |  |  |  | 1 |
| *Pica pica* |  |  |  |  |  |  |  |  |  |  |  |  | 3 | 2 |  |  |  |  |  | 5 |
| *Saxicola rubicola* |  |  |  |  |  | 2 |  |  |  |  |  |  |  |  | 5 |  |  |  |  | 7 |
| *Serinus serinus* |  |  |  |  |  |  |  |  |  |  |  |  | 1 |  |  |  |  |  |  | 1 |
| *Sitta europaea* |  |  |  |  |  |  |  | 2 |  |  |  |  |  |  | 1 |  |  |  |  | 3 |
| *Streptopelia decaocto* |  | 1 | 1 | 0 | 4 | 15 | 2 | 2 | 13 |  | 3 | 2 | 8 | 8 | 7 |  | 10 | 1 |  | 77 |
| *Streptopelia turtur* |  |  |  |  |  |  |  | 3 |  |  |  |  |  |  | 6 | 2 |  |  |  | 11 |
| *Strix aluco* |  |  |  |  |  |  |  |  |  |  |  |  |  |  | 2 |  |  |  |  | 2 |
| *Sturnus unicolor* |  |  | 1 |  |  |  |  | 1 |  |  |  |  |  |  | 4 |  |  |  |  | 6 |
| *Sylvia atricapilla* |  |  |  |  |  | 1 |  | 3 |  |  |  |  |  |  |  |  |  |  |  | 4 |
| *Sylvia borin* |  |  |  |  |  | 1 |  |  |  |  |  |  |  |  |  |  |  |  |  | 1 |
| *Curruca iberiae* |  |  |  |  |  |  |  |  |  |  |  |  |  |  | 3 | 1 |  |  |  | 4 |
| *Curruca communis* |  | 1 |  |  |  |  |  | 1 |  |  |  |  |  |  | 1 |  |  |  |  | 3 |
| *Curruca melanocephala* |  |  |  |  |  | 1 |  | 4 |  |  |  |  | 4 |  | 11 | 1 |  |  |  | 21 |
| *Tringa ochropus* |  |  |  |  |  |  |  |  |  |  |  |  |  |  | 2 |  |  |  |  | 2 |
| *Troglodytes troglodytes* |  |  |  |  |  |  |  |  |  |  |  |  |  |  | 1 |  |  |  |  | 1 |
| *Turdus merula* |  |  |  |  | 1 | 9 |  | 3 |  |  |  |  | 2 |  | 9 |  |  |  |  | 24 |
| *Turdus philomelos* |  |  | 1 |  |  | 1 |  |  |  |  |  |  |  |  |  |  |  |  |  | 2 |
| *Upupa epops* |  |  |  |  |  |  |  |  |  |  |  |  |  |  | 3 |  |  |  |  | 3 |
| Total | 4 | 91 | 61 | 78 | 6 | 92 | 38 | 134 | 94 | 79 | 65 | 2 | 112 | 78 | 205 | 15 | 109 | 60 | 44 | 1367 |
|  | 156 |  |  | 84 |  | 92 | 172 |  | 173 |  | 67 |  | 112 | 283 |  | 228 |  |  |  | 1367 |

**Table S2.** Seroprevalences (by ELISA and VNT) and prevalences (by RT-qPCR) obtained for the complete sample set encompassing all bird species from September/October 2021 to December 2023. West Nile virus (WNV), Usutu virus (USUV), Bagaza virus (BAGV) and unspecified antibodies (Undetermined flavivirus). Serological results for *A. rufa* from 2021-2022 were previously reported in Fontoura-Gonçalves et al. (2025) and are included here for completeness and temporal comparison; all remaining results including the RT-qPCR for 2021-2023 and serological results from 2022 onward and all the data for other bird species are original to the present study. The number of samples analysed for USUV was lower than for WNV and BAGV because in some individuals insufficient serum volume remained to test at USUV VNT dilutions >1:20. Values are presented as % (number positive/total number analysed).

| Year | 2021 |  | 2022 |  |  |  |  | 2023 |  |  |  |  |  | Total<br>(%) |
| --- | --- | --- | --- | --- | --- | --- | --- | --- | --- | --- | --- | --- | --- | --- |
| Season | Autumn |  | Winter | Spring | Summer |  | Autumn | Winter | Spring | Summer |  | Autumn |  |  |
| Month | Sep/Oct | Nov | Feb | Apr | Jun | Aug | Oct/Nov | Feb | Apr | Jun | Aug | Sep/Oct | Nov/Dec |  |
| ELISA<br>Seroprevalence<br>(%) | 66.67<br>(34/51) | 67.39<br>(31/46) | 73.33<br>(33/45) | 42.55<br>(40/94) | 62.16<br>(23/37) | 30.08<br>(37/123) | 61.29<br>(38/62) | 55.77<br>(29/52) | 44.54<br>(53/119) | 62.34<br>(48/77) | 38.76<br>(69/178) | 34.48<br>(40/116) | 35<br>(21/60) | 46.36<br>(496/1070) |
|  | 67.01 (65/97) |  | 37.50 (60/160) |  |  |  |  | 45.88 (117/255) |  |  |  |  | 34.66 (61/176) |  |
| Seroprevalence by VNT (%): |  |  |  |  |  |  |  |  |  |  |  |  |  |  |
| WNV | 27.08<br>(13/48) | 27.50<br>(11/40) | 28.89<br>(13/45) | 30.38<br>(24/79) | 22.86<br>(8/35) | 23.76<br>(24/101) | 35.71<br>(20/56) | 31.37<br>(16/51) | 44.71<br>(38/85) | 31.58<br>(24/76) | 22.29<br>(35/157) | 25.27<br>(23/91) | 21.74<br>(10/46) | 28.46<br>(259/910) |
|  | 4.17<br>(2/48) | 2.56<br>(1/39) | 0<br>(0/45) | 0<br>(0/51) | 0<br>(0/35) | 1.54<br>(1/65) | 0<br>(0/49) | 0<br>(0/38) | 0<br>(0/51) | 1.92<br>(1/52) | 3.85<br>(3/78) | 0<br>(0/47) | 0<br>(0/25) | 1.28<br>(8/623) |
| BAGV | 8.33<br>(4/48) | 25<br>(10/40) | 15.56<br>(7/45) | 6.33<br>(5/79) | 20<br>(7/35) | 0.99<br>(1/101) | 1.79<br>(1/56) | 13.73<br>(7/51) | 2.35<br>(2/85) | 7.89<br>(6/76) | 1.91<br>(3/157) | 4.40<br>(4/91) | 6.52<br>(3/46) | 6.59<br>(60/910) |
|  | 35.42<br>(17/48) | 30<br>(12/40) | 37.78<br>(17/45) | 17.72<br>(14/79) | 31.43<br>(11/35) | 6.93<br>(7/101) | 25<br>(14/56) | 17.65<br>(9/51) | 8.24<br>(7/85) | 23.68<br>(18/76) | 13.38<br>(21/157) | 18.68<br>(17/91) | 26.09<br>(12/46) | 19.34<br>(176/910) |
| Total VNT | 75<br>(36/48) | 85<br>(34/40) | 82.22<br>(37/45) | 54.43<br>(43/79) | 74.29<br>(26/35) | 32.67<br>(33/101) | 62.50<br>(35/56) | 62.75<br>(32/51) | 55.29<br>(47/85) | 64.47<br>(49/76) | 40.13<br>(63/157) | 48.35<br>(44/91) | 54.35<br>(25/46) | 55.27<br>(503/910) |
|  | 79.55 (70/88) |  | 43.38 (59/136) |  |  |  |  | 48.07 (112/233) |  |  |  |  | 50.36 (69/137) |  |
| Prevalences by RT-qPCR (%): |  |  |  |  |  |  |  |  |  |  |  |  |  |  |
| WNV | 2.17<br>(2/92) | 0<br>(0/60) | 0<br>(0/78) | 0<br>(0/58) | 0<br>(0/35) | 1.79<br>(2/112) | 4.09<br>(7/171) | 0<br>(0/60) | 0<br>(0/38) | 0<br>(0/73) | 1.76<br>(3/170) | 0.83<br>(1/121) | 2.91<br>(3/103) | 1.54<br>(18/1171) |
|  | 0<br>(0/92) | 1.67<br>(1/60) | 0<br>(0/78) | 0<br>(0/58) | 0<br>(0/35) | 0<br>(0/112) | 0<br>(0/171) | 0<br>(0/60) | 0<br>(0/38) | 0<br>(0/73) | 0<br>(0/170) | 0<br>(0/121) | 0.97<br>(1/103) | 0.17<br>(2/1171) |
| BAGV | 19.57<br>(18/92) | 5<br>(3/60) | 0<br>(0/78) | 0<br>(0/58) | 0<br>(0/35) | 0<br>(0/112) | 0<br>(0/171) | 0<br>(0/60) | 0<br>(0/38) | 0<br>(0/73) | 0<br>(0/170) | 0<br>(0/121) | 0<br>(0/1039) | 1.79<br>(21/1171) |
|  | 21.74<br>(20/92) | 6.67<br>(4/60) | 0<br>(0/78) | 0<br>(0/58) | 0<br>(0/35) | 0<br>(0/112) | 4.09<br>(7/171) | 0<br>(0/60) | 0<br>(0/38) | 0<br>(0/73) | 1.76<br>(3/170) | 0.83<br>(1/121) | 3.88<br>(4/103) | 3.50<br>(41/1171) |
| 15.79 (24/152) |  | 0 (0/147) |  |  |  |  | 1.23 (3/243) |  |  |  |  | 2.23 | 5/224) |  |

**Table S3.** Seroprevalences (by ELISA and VNT) and prevalences (by RT-qPCR) for red-legged partridges (*A. rufa*) sampled every two months from September/October 2021 to December 2023 in Serpa (A) and in Autumn 2023 in Mértola (B). West Nile virus (WNV), Usutu virus (USUV), Bagaza virus (BAGV) and unspecified antibodies (Undetermined flavivirus). Serological results for *A. rufa* from 2021-2022 were previously reported in Fontoura-Gonçalves et al. (2025) and are included here for completeness and temporal comparison; all remaining results including the RT-qPCRs for 2021-2022 and serological results from 2022 onward are original to the present study. The number of samples analysed for USUV was lower than for WNV and BAGV because in some individuals insufficient serum volume remained to test at USUV VNT dilutions >1:20. Values are presented as % (number positive/total number analysed).

A. Serpa

| Year | 2021 |  | 2022 |  |  |  |  | 2023 |  |  |  |  | Total<br>(%) |  |
| --- | --- | --- | --- | --- | --- | --- | --- | --- | --- | --- | --- | --- | --- | --- |
| Season | Autumn |  | Winter | Spring | Summer |  | Autumn | Winter | Spring | Summer |  | Autumn |  |  |
| Month | Sep/Oct | Nov | Feb | Apr | Jun | Aug | Oct/Nov | Feb | Apr | Jun | Aug | Sep/Oct |  | Nov/Dec |
| ELISA Seroprevalence (%) | 66.67<br>(34/51) | 70<br>(28/40) | 73.33<br>(33/45) | 64.29<br>(18/28) | 62.86<br>(22/35) | 16.28<br>(7/43) | 68.75<br>(33/48) | 55.77<br>(29/52) | 70.27<br>(26/37) | 72.92<br>(35/48) | 74.51<br>(38/51) | 63.64<br>(35/55) | 51.22<br>(21/41) | 62.54<br>(359/574) |
| Seroprevalence by VNT (%): |  |  |  |  |  |  |  |  |  |  |  |  |  |  |
| WNV | 27.08<br>(13/48) | 26.32<br>(10/38) | 28.89<br>(13/45) | 35.71<br>(10/28) | 21.88<br>(7/32) | 9.52<br>(4/42) | 36.36<br>(16/44) | 31.37<br>(16/51) | 52.94<br>(18/34) | 34.04<br>(16/47) | 29.41<br>(15/51) | 51.35<br>(19/37) | 37.04<br>(10/27) | 31.87<br>(167/524) |
| USUV | 4.17<br>(2/48) | 2.63<br>(1/38) | 0<br>(0/45) | 0<br>(0/28) | 0<br>(0/32) | 0<br>(0/42) | 0<br>(0/44) | 0<br>(0/38) | 0<br>(0/29) | 0<br>(0/38) | 2.56<br>(1/39) | 0<br>(0/34) | 0<br>(0/21) | 0.84<br>(4/476) |
| BAGV | 8.33<br>(4/48) | 26.32<br>(10/38) | 15.56<br>(7/45) | 17.86<br>(5/28) | 21.88<br>(7/32) | 2.38<br>(1/42) | 2.27<br>(1/44) | 13.73<br>(7/51) | 5.88<br>(2/34) | 10.64<br>(5/47) | 5.88<br>(3/51) | 10.81<br>(4/37) | 11.11<br>(3/27) | 11.26<br>(59/524) |
| Undetermined flaviviruses | 35.42<br>(17/48) | 31.58<br>(12/38) | 37.78<br>(17/45) | 32.14<br>(9/28) | 34.38<br>(11/32) | 11.90<br>(5/42) | 31.82<br>(14/44) | 17.65<br>(9/51) | 14.71<br>(5/34) | 29.79<br>(14/47) | 23.53<br>(12/51) | 32.43<br>(12/37) | 33.33<br>(9/27) | 27.86<br>(146/524) |
| Total VNT | 75<br>(36/48) | 86.84<br>(33/38) | 82.22<br>(37/45) | 85.71<br>(24/28) | 78.13<br>(25/32) | 23.81<br>(10/42) | 70.45<br>(31/44) | 62.75<br>(32/51) | 73.53<br>(25/34) | 74.47<br>(35/47) | 60.78<br>(31/51) | 94.59<br>(35/37) | 81.48<br>(22/27) | 71.76<br>(375/524) |
|  | 80.23 (69/86) |  | 47.30 (35/74) |  |  |  |  | 67.35 (66/98) |  |  |  |  | 89.06 (57/64) |  |
| Prevalence by RT-qPCR (%): |  |  |  |  |  |  |  |  |  |  |  |  |  |  |
| WNV | 2.17<br>(2/92) | 0<br>(0/58) | 0<br>(0/78) | 0<br>(0/26) | 0<br>(0/33) | 0<br>(0/42) | 4.4<br>(7/159) | 0<br>(0/60) | 0<br>(0/32) | 0<br>(0/48) | 3.92<br>(2/51) | 0<br>(0/61) | 4.11<br>(3/73) | 1.72<br>(14/813) |
| USUV | 0<br>(0/92) | 1.72<br>(1/58) | 0<br>(0/78) | 0<br>(0/26) | 0<br>(0/33) | 0<br>(0/42) | 0<br>(0/159) | 0<br>(0/60) | 0<br>(0/32) | 0<br>(0/48) | 0<br>(0/51) | 0<br>(0/61) | 0<br>(0/73) | 0.12<br>(1/813) |
| BAGV | 19.57<br>(18/92) | 3.45<br>(2/58) | 0<br>(0/78) | 0<br>(0/26) | 0<br>(0/33) | 0<br>(0/42) | 0<br>(0/159) | 0<br>(0/60) | 0<br>(0/32) | 0<br>(0/48) | 0<br>(0/51) | 0<br>(0/61) | 0<br>(0/73) | 2.46<br>(20/813) |
| Total Orthoflavivirus prevalence | 21.74<br>(20/92) | 5.17<br>(3/58) | 0<br>(0/78) | 0<br>(0/26) | 0<br>(0/33) | 0<br>(0/42) | 4.4<br>(7/159) | 0<br>(0/60) | 0<br>(0/32) | 0<br>(0/48) | 3.92<br>(2/51) | 0<br>(0/61) | 4.11<br>(3/73) | 4.3<br>(35/813) |
|  | 15.33 (23/150) |  | 0 (0/75) |  |  |  |  | 2.02 (2/99) |  |  |  |  | 2.24 (3/134) |  |

B. Mértola

| Year | 2023 |  | Total (%) |
| --- | --- | --- | --- |
| Season | Autumn |  |  |
| Month | Sep/Oct | Nov/Dec |  |
| ELISA Seroprevalence (%) | 5<br>(2/40) | 0<br>(0/19) | 2.89<br>(2/69) |
| Seroprevalence by VNT (%): |  |  |  |
| WNV | 5.89<br>(2/34) | 0<br>(0/19) | 3.77<br>(2/53) |
| USUV | 0<br>(0/8) | 0<br>(0/4) | 0<br>(0/12) |
| BAGV | 0<br>(0/34) | 0<br>(0/19) | 0<br>(0/53) |
| Undetermined flaviviruses | 8.82<br>(3/34) | 15.79<br>(3/19) | 11.32<br>(6/53) |
| Total VNT | 14.71<br>(5/34) | 15.79<br>(3/19) | 15.09<br>(8/53) |
| Prevalence by RT-qPCR (%): |  |  |  |
| WNV | 2.5<br>(1/40) | 0<br>(0/30) | 1.43<br>(1/70) |
| USUV | 0<br>(0/40) | 3.33<br>(1/30) | 1.43<br>(1/70) |
| BAGV | 0<br>(0/40) | 0<br>(0/30) | 0<br>(0/70) |
| Total Orthoflavivirus prevalence | 2.5<br>(1/40) | 3.33<br>(1/30) | 2.86<br>(2/70) |

**Table S4.** Mosquitoes captured in Serpa and Mértola and morphologically identified by sex and date. Only female specimens were taxonomically identified by genera, species or species/complex. Values are presented as % (number positive/total number analysed).

| Year | 2022 |  | 2023 |  |  |  |  | Total | %<br>(n Captured<br>/Total of<br>Females) |
| --- | --- | --- | --- | --- | --- | --- | --- | --- | --- |
| Season | Summer | Autumn | Winter | Spring | Summer |  | Autumn |  |  |
| Month | Aug | Oct | Feb | Apr | Jun | Aug | Oct |  |  |
| Females: | 16 | 18 | 4 | 25 | 51 | 16 | 70 | 200 |  |
| **Culex s.l.** | **14** | **13** | **3** | **18** | **43** | **16** | **67** | **174** | **87.0% (174/200)** |
| *Culex pipiens/torrentium* | 2 | 4 |  | 2 | 16 | 1 | 9 | 34 | 17.0% (34/200) |
| *Culex univittatus/perexiguus* | 7<br>(1WNV positive) | 3<br>(1 WNV positive) |  | 4 | 5 | 6 | 8 | 33 | 16.50% (33/200) |
| *Culex theileri* | 2 |  |  | 4 | 4 | 2 | 6 | 18 | 9% (18/200) |
| *Culex mimeticus* |  | 1 |  | 2 | 3 |  |  | 6 | 3% (6/200) |
| *Culex hortensis* |  |  |  |  |  |  | 1 | 1 | 0.50% (1/200) |
| *Culex* sp.. | 3 | 5 | 3 | 6 | 15 | 7 | 43 | 82 | 4% (82/200) |
| **Aedes s.l.** | **2** | **4** |  | **2** | **3** |  |  | **11** | **5.50% (11/200)** |
| *Aedes berlandi* | 2 | 2 |  | 2 | 2 |  |  | 8 | 4% (8/200) |
| *Aedes finlaya/genitaculus* |  | 1 |  |  |  |  |  | 1 | 0.50% (1/200) |
| *Aedes* sp. |  | 1 |  |  | 1 |  |  | 2 | 1% (2/100) |
| **Anopheles s.l.** |  | **1** |  | **4** | **3** |  | **2** | **10** | **5% (10/200)** |
| *Anopheles maculopenis*<br>complex |  |  |  | 1 | 3 |  |  | 4 | 2% (4/200) |
| *Anopheles* sp. |  | 1 |  | 3 |  |  | 2 | 6 | 3% (6/200) |
| **Culiseta s.l.** |  |  | **1** | **1** | **2** |  | **1** | **5** | **2.50 (5/200)** |
| *Culiseta annulata* |  |  | 1 |  |  |  |  | 1 | 0.50% (1/200) |
| *Culiseta longireolata* |  |  |  |  | 1 |  |  | 1 | 0.50% (1/200) |
| *Culiseta* sp. |  |  |  | 1 | 1 |  |  | 2 | 1% (2/200) |
| *Culiseta litorea* |  |  |  |  |  |  | 1 | 1 | 0.50% (1/200) |
| Males | 4 | 6 | 1 | 5 | 9 | 0 | 25 | 50 | - |
| No possible species or sex ID | 0 | 1 | 0 | 0 | 4 | 0 | 0 | 5 | - |
| **Total mosquitos** | **20** | **25** | **5** | **30** | **64** | **16** | **95** | **255** | **-** |

**Table S5.** Blood-fed females by capture date. Blood-fed females were identified morphologically and further confirmed by COI sequencing. For four specimens, for which molecular identification was inconclusive, the species reported here is based on morphological identification. Values are presented as % (number positive/total number analysed). COI sequences obtained were deposited in GenBank, accession numbers: PZ338730-44; PZ364570-71; PZ383187- PZ383200; PZ385223- PZ385247.

| Year | 2022 |  | 2023 |  |  |  | Total | % (n species/total blood-fed females) |
| --- | --- | --- | --- | --- | --- | --- | --- | --- |
| Month | Aug | Oct | Apr | Jun | Aug | Oct |  |  |
| **Total Culex spp.:** | **6** | **3** | **3** | **9** | **3** | **28** | **52** | **88.14% (52/59)** |
| **Culex pipiens\*:** | **2** | **1** | **1** | **5** | **2** | **16** | **27** | **45.76% (27/59)** |
| *Culex pipiens*\* | 2 | 1 | 1 | 4 | 1 | 15 | 24 | 40.68% (24/59) |
| *Culex pipiens molestus*\* |  |  |  | 1 |  | 1 | 2 | 3.39% (2/59) |
| *Culex pipiens pipiens*\* |  |  |  |  | 1 |  | 1 | 1.69% (1/59) |
| *Culex theileri*\* |  |  | 1 | 1 |  | 8 | 10 | 16.95% (10/59) |
| *Culex univittatus*\* | 4 | 1 |  | 2 | 1 | 1 | 9 | 15.25% (9/59) |
| *Culex mimeticus*\* |  | 1 | 1 |  |  |  | 2 | 3.39% (2/59) |
| *Culex laticinctus*\* |  |  |  | 1 |  |  | 1 | 1.69% (1/59) |
| *Culex pipiens/torrentium* |  |  |  |  |  | 1 | 1 | 1.69% (1/59) |
| *Culex univittatus/perexiguus* |  |  |  |  |  | 2 | 2 | 3.39% (1/59) |
| **Total Aedes spp.** | **0** | **1** | **0** | **2** | **0** | **0** | **3** | **5.08% (3/59)** |
| *Aedes caspius*\* |  |  |  | 2 |  |  | 2 | 3.39% (2/59) |
| *Aedes geniculatus*\* |  | 1 |  |  |  |  | 1 | 1.69% (1/59) |
| **Total Culiseta spp.** | **0** | **0** | **0** | **2** | **0** | **1** | **3** | **5.08% (3/59)** |
| *Culiseta longireolata*\* |  |  |  | 2 |  |  | 2 | 3.39% (2/59) |
| *Culiseta sp (litorea)* |  |  |  |  |  | 1 | 1 | 1.69% (1/59) |
| *Anopheles atroparvus*\* |  |  |  | 1 |  |  | 1 | 1.69% (1/59) |
| **Total** | **6** | **4** | **3** | **14** | **3** | **29** | **59** |  |

(i)\*molecular identification confirmed.

**Table S6.** West Nile Virus (WNV) molecular detections (by RT-qPCR) and specific antibodies (by VNT) by bird species sampled from Autumn 2021 to Summer 2023. Values are presented as % (number positive/total number analysed).

| WNV |  |  |  |  |  |  |  |  |  |  |  |  |
| --- | --- | --- | --- | --- | --- | --- | --- | --- | --- | --- | --- | --- |
| Year | 2021 |  | 2022 |  |  |  | 2023 |  |  |  | Total % |  |
| Season | Autumn |  | Spring |  | Summer/Autumn |  | Spring |  | Summer |  |  |  |
|  | PCR | VNT | PCR | VNT | PCR | VNT | PCR | VNT | PCR | VNT | PCR | VNT |
| *Coccothraustes coccothraustes*\* |  |  |  |  | 0<br>(0/3) | 100<br>(2/2) |  | 76.19<br>(16/21) | 0<br>(0/2) | 100<br>(1/1) | 0<br>(0/5) | 83.33<br>(20/24) |
| *Columba livia*\* |  |  | 0<br>(0/10) | 33.33<br>(4/12) | 0<br>(0/12) | 38.46<br>(5/13) |  |  | 0<br>(0/13) | 41.67<br>(5/13) | 0<br>(0/35) | 35.9<br>(14/38) |
| *Streptopelia decaocto*\* |  | 50<br>(1/2) | 0<br>(0/15) | 26.67<br>(4/15) | 0<br>(0/13) | 40<br>(6/15) | 0<br>(0/3) | 14.29<br>(1/7) | 0<br>(0/23) | 20<br>(5/25) | 0<br>(0/54) | 25<br>(17/65) |
| ***Cyanopica cooki*\*** |  |  |  | 25<br>(1/4) | **9.09**<br>**(1/11)** | 20<br>(2/10) |  | 100<br>(1/1) | **4.17**<br>**(1/25)** | 10.53<br>(2/20) | 5.56<br>(2/36) | 17.14<br>(6/35) |
| *Turdus merula*\* |  |  |  | 44.44<br>(4/9) | 0<br>(0/3) | 33.33<br>(1/3) |  | 0<br>(0/2) | 0<br>(0/8) | 11.11<br>(1/9) | 0<br>(0/11) | 26.09<br>(6/23) |
| *Actitis hypoleucos* |  |  |  |  |  |  |  |  |  | 100<br>(1/1) | 0<br>(0/0) | 100<br>(1/1) |
| *Columba palumbus* |  |  |  |  | 0<br>(0/1) | 100<br>(1/1) |  |  | 0<br>(0/12) | 16.67<br>(2/12) | 0<br>(0/13) | 23.08<br>(3/13) |
| *Dendrocopos major* |  |  |  |  | 0<br>(0/1) | 0<br>(0/1) |  | 0<br>(0/1) | 0<br>(0/5) | 33.33<br>(2/6) | 0<br>(0/6) | 25<br>(2/8) |
| *Fringilla coelebs* |  |  | 0<br>(0/1) |  | 0<br>(0/3) | 33.33<br>(1/3) |  |  | 0<br>(0/2) | 0<br>(0/2) | 0<br>(0/6) | 20<br>(1/5) |
| ***Garrulus glandarius*** |  |  |  | 100<br>(1/1) | **33.33**<br>**(1/3)** | 66.67<br>(2/3) |  | 100<br>(1/1) | 0<br>(0/3) | 75<br>(3/4) | 6.25<br>(1/6) | 77.78<br>(7/9) |
| *Merops apiaster* |  |  |  |  | 0<br>(0/1) | 33.33<br>(1/3) |  |  |  |  | 0<br>(0/1) | 33.33<br>(1/3) |
| *Oriolus oriolus* |  |  |  |  | 0<br>(0/1) | 100<br>(1/1) |  |  |  |  | 0<br>(0/1) | 100<br>(1/1) |
| *Parus major* |  |  |  |  | 0<br>(0/5) | 50<br>(1/2) |  |  | 0<br>(0/7) | 33.33<br>(2/6) | 0<br>(0/12) | 37.5<br>(3/8) |
| *Pica pica* |  |  |  |  |  |  |  | 33.33<br>(1/3) | 0<br>(0/2) | 0<br>(0/2) | 0<br>(0/2) | 20<br>(1/5) |

**Table S6** (continued). West Nile Virus (WNV) molecular detections (by RT-qPCR) and specific antibodies (by VNT) by bird species sampled from Autumn 2021 to Summer 2023. Values are presented as % (number positive/total number analysed).

| WNV |  |  |  |  |  |  |  |  |  |  |  |  |
| --- | --- | --- | --- | --- | --- | --- | --- | --- | --- | --- | --- | --- |
| Year | 2021 |  | 2022 |  |  |  | 2023 |  |  |  | Total % |  |
| Season | Autumn |  | Spring |  | Summer/Autumn |  | Spring |  | Summer |  |  |  |
|  | PCR | VNT | PCR | VNT | PCR | VNT | PCR | VNT | PCR | VNT | PCR | VNT |
| *Sitta europaea* |  |  |  |  | 0<br>(0/2) | 50<br>(1/2) |  |  | 0<br>(0/1) |  | 0<br>(0/3) | 50<br>(1/2) |
| *Streptopelia turtur* |  |  |  |  | 0<br>(0/3) | 0<br>(0/2) |  |  | 0<br>(0/8) | 37.5<br>(3/8) | 0<br>(0/11) | 30<br>(3/10) |
| *Strix aluco* |  |  |  |  |  |  |  |  | 0<br>(0/2) | 100<br>(2/2) | 0<br>(0/2) | 100<br>(2/2) |
| *Sturnus unicolor* |  |  |  |  | 0<br>(0/1) |  |  |  | 0<br>(0/4) | 25<br>(1/4) | 0<br>(0/5) | 20<br>(1/5) |
| Other species (negative) | 1 | 0 | 6 | 10 | 33 | 13 | 3 | 14 | 73 | 40 | 116 | 77 |
| Total | 0<br>(0/1) | 50<br>(1/2) | 0<br>(0/26) | 27.45<br>(14/51) | 2.11<br>(2/95) | 32.88<br>(24/73) | 0<br>(0/3) | 39.22<br>(20/51) | 0.55<br>(1/183) | 19.35<br>(30/155) | 0.91<br>(3/326) | 26.41<br>(89/333) |

(i) In bold are the species where molecular detection by RT-qPCR was confirmed. \*Species that present a total of analysed individuals higher than 20.

**Table S7.** Seroprevalences (by ELISA and VNT) and prevalences (by RT-qPCR) by bird species (excluding *A. rufa*) sampled in Autumn 2021 and Spring and Summer/Autumn in 2022 and 2023 in Serpa (A) and in Summer 2023 in Mértola (B). West Nile virus (WNV), Usutu virus (USUV), Bagaza virus (BAGV) and unspecified antibodies (Undetermined flavivirus). The number of samples analysed for USUV was lower than for WNV and BAGV because in some individuals insufficient serum volume remained to test at USUV VNT dilutions >1:20. Values are presented as % (number positive/total number analysed).

A. Serpa

| Year | 2021 | 2022 |  |  |  | 2023 |  |  |  | Total % |
| --- | --- | --- | --- | --- | --- | --- | --- | --- | --- | --- |
| Season | Autumn | Spring | Summer/Autumn |  |  | Spring | Summer/Autumn |  |  |  |
| Month | Oct/Nov | Apr | Jun | Aug | Oct | Apr | Jun | Aug | Sep/Oct |  |
| ELISA Seroprevalence (%) | 50<br>(3/6) | 33.33<br>(22/66) | 50<br>(1/2) | 37.5<br>(30/80) | 35.71<br>(5/14) | 32.93<br>(27/82) | 44.83<br>(13/29) | 34.25<br>(25/73) | 14.29<br>(3/21) | 34.58<br>(129/373) |
|  |  |  | 37.5 (36/96) |  |  |  | 33.33 (41/123) |  |  |  |
| Seroprevalence by VNT (%) |  |  |  |  |  |  |  |  |  |  |
| WNV | 50<br>(1/2) | 27.45<br>(14/51) | 33.33<br>(1/3) | 33.9<br>(20/59) | 33.33<br>(4/12) | 39.22<br>(20/51) | 27.59<br>(8/29) | 28.13<br>(18/64) | 10<br>(2/20) | 30.24<br>(88/291) |
|  |  |  | 33.78(25/74) |  |  |  | 24.78 (28/113) |  |  |  |
| USUV | 0<br>(0/1) | 0<br>(0/23) | 0<br>(0/3) | 4.35<br>(1/23) | 0<br>(0/5) | 0<br>(0/22) | 7.14<br>(1/14) | 0<br>(0/22) | 0<br>(0/5) | 1.69<br>(2/118) |
|  |  |  | 3.23 (1/31) |  |  |  | 2.44 (1/41) |  |  |  |
| BAGV | 0<br>(0/2) | 0<br>(0/51) |  | 0<br>(0/3) |  | 0<br>(0/59) |  | 0<br>(0/12) |  | 0<br>(0/51) |
|  |  |  | 0 (0/74) |  |  |  | 0.88 (1/ 113) |  |  |  |
| Undetermined flaviviruses | 0<br>(0/2) | 9.8<br>(5/51) | 0<br>(0/3) | 3.39<br>(2/59) | 0<br>(0/12) | 3.92<br>(2/51) | 13.79<br>(4/29) | 9.38<br>(6/64) | 10<br>(2/20) | 7.22<br>(21/291) |
|  |  |  | 2.70 (2/74) |  |  |  | 10.62 (12/113) |  |  |  |
| Total VNT | 50<br>(1/2) | 37.25<br>(19/51) | 33.33<br>(1/3) | 38.98<br>(23/59) | 33.33<br>(4/12) | 43.14<br>(22/51) | 48.28<br>(14/29) | 37.5<br>(24/64) | 20<br>(4/20) | 38.49<br>(112/291) |
|  |  |  | 37.84 (28/74) |  |  |  | 37.17 (42/113) |  |  |  |
| Prevalences by RT-qPCR (%): |  |  |  |  |  |  |  |  |  |  |
| WNV | 0<br>(0/2) | 0<br>(0/32) | 0<br>(0/2) | 2.86<br>(2/70) | 0<br>(0/12) | 0<br>(0/6) | 0<br>(0/25) | 0<br>(0/49) | 0<br>(0/20) | 0.92<br>(2/218) |
| USUV | 0<br>(0/2) | 0<br>(0/32) | 0<br>(0/2) | 0<br>(0/70) | 0<br>(0/12) | 0<br>(0/6) | 0<br>(0/25) | 0<br>(0/49) | 0<br>(0/20) | 0<br>(0/218) |
| BAGV | 50<br>(1/2) | 0<br>(0/32) | 0<br>(0/2) | 0<br>(0/70) | 0<br>(0/12) | 0<br>(0/6) | 0<br>(0/25) | 0<br>(0/49) | 0<br>(0/20) | 0.46<br>(1/218) |
| Total prevalence by RT-qPCR | 50<br>(1/2) | 0<br>(0/32) | 0<br>(0/2) | 2.86<br>(2/70) | 0<br>(0/12) | 0<br>(0/6) | 0<br>(0/25) | 0<br>(0/49) | 0<br>(0/20) | 1.38<br>(3/218) |
|  |  |  | 2.38 (2/72) |  |  |  | 0 (0/94) |  |  |  |

B. Mértola

|  |  |
| --- | --- |
| **Year** | **2023** |
| **Season** | **Summer** |
| **Month** | **Aug** |
| **ELISA Seroprevalence (%)** | 11.11<br>(6/54) |
| **Seroprevalence by VNT (%)** |  |
| **WNV** | 4.76<br>(2/42) |
| **USUV** | 5.88<br>(1/17) |
| **BAGV** | 0<br>(0/42) |
| **Undetermined flaviviruses** | 7.14<br>(3/42) |
| **Total VNT** | 19.05<br>(8/42) |
| **Prevalences by RT-qPCR (%):** |  |
| **WNV** | 1.43<br>(1/70) |
| **USUV** | 0<br>(0/70) |
| **BAGV** | 0<br>(0/70) |
| **Total prevalence by RT-qPCR** | 1.43<br>(1/70) |

**Table S8.** Full and partial genomes obtained in this study. Blue for West Nile Virus (WNV) sequences, green for Usutu Virus (USUV) sequences and orange for Bagaza Virus (BAGV) sequences.

| GenBank accession | COD | Virus | Year | Month | Species | Location | Total reads | Mapped reads | Genome coverage % | Nº con tigs | Total bp | Mean coverage | Min. coverage |
| --- | --- | --- | --- | --- | --- | --- | --- | --- | --- | --- | --- | --- | --- |
| PZ377065 | Av2021_065 | WNV | 2021 | Oct | *Alectoris rufa* | Serpa | 32608324 | 948061 | 99.9 | 1 | 11030 | 14478.0 | 20 |
| PZ377064 | LV22672 | WNV | 2022 | Oct | *Alectoris rufa* | Serpa | 3219756 | 84934 | 100.0 | 1 | 11030 | 1386.0 | 2 |
| PZ377063 | LV23201 | WNV | 2023 | Aug | *Alectoris rufa* | Serpa | 36447664 | 2982353 | 100.0 | 1 | 11030 | 46273.0 | 2 |
| PZ377061 | Av2023_055 | WNV | 2023 | Oct | *Alectoris rufa* | Mértola | 4129292 | 14125 | 99.8 | 1 | 11001 | 210.3 | 2 |
| PZ377062 | F67551 | WNV | 2023 | Aug | *Cyanopica cooki* | Mértola | 3077792 | 1206 | 93.7 | 1 | 10332 | 16.4 | 2 |
| PZ392167-PZ392171 | F67652 | WNV | 2022 | Aug | *Cyanopica cooki* | Mértola | 2779382 | 24 | 7.5 | 5 | 822 | 3.3 | 2 |
| PZ400790-PZ400794 | Gaio 1A | WNV | 2022 | Aug | *Garrulus glandarius* | Serpa | 39578310 | 33 | 7.0 | 5 | 775 | 5.5 | 2 |
| PZ377060 | 22A | WNV | 2022 | Oct | *Culex univitattus* | Serpa | 31863952 | 71037 | 99.6 | 1 | 10984 | 1148.0 | 73 |
| PZ400796-PZ400817 | 60A | WNV | 2022 | Aug | *Culex univitattus* | Serpa | 31386078 | 1356 | 68.6 | 22 | 7563 | 26.5 | 2 |
| PQ677887 | Romeiras penas 219 | USUV | 2023 | Nov | *Alectoris rufa* | Mértola | 29220938 | 7,550 | 100.0 | 1 | 11108 | 105.0 | 14 |
| PQ672170-79 | Av2021_061 | USUV | 2021 | Nov | *Alectoris rufa* | Serpa | 558636 | 113 | 12.6 | 10 | 1401 | 5.0 | 1 |
| PZ377066 | Av2021_008 | BAGV | 2021 | Oct | *Alectoris rufa* | Serpa | 9517340 | 1799088 | 100.0 | 1 | 10944 | 309617.0 | 31 |
| PZ392148-PZ392165 | Av2021_001 | BAGV | 2021 | Oct | *Emberiza calandra* | Serpa | 32128352 | 110 | 38.2 | 23 | 4178 | 4.0 | 2 |

(i) USUV genome and partial sequences (accession numbers: PQ677887; PQ672170–PQ672179) have been previously characterized as belonging to the African 3.1 sub-lineage (Queirós et al., 2025). These detections were included in the present study for an overall analysis of orthoflaviviruses circulation and genomic diversity.

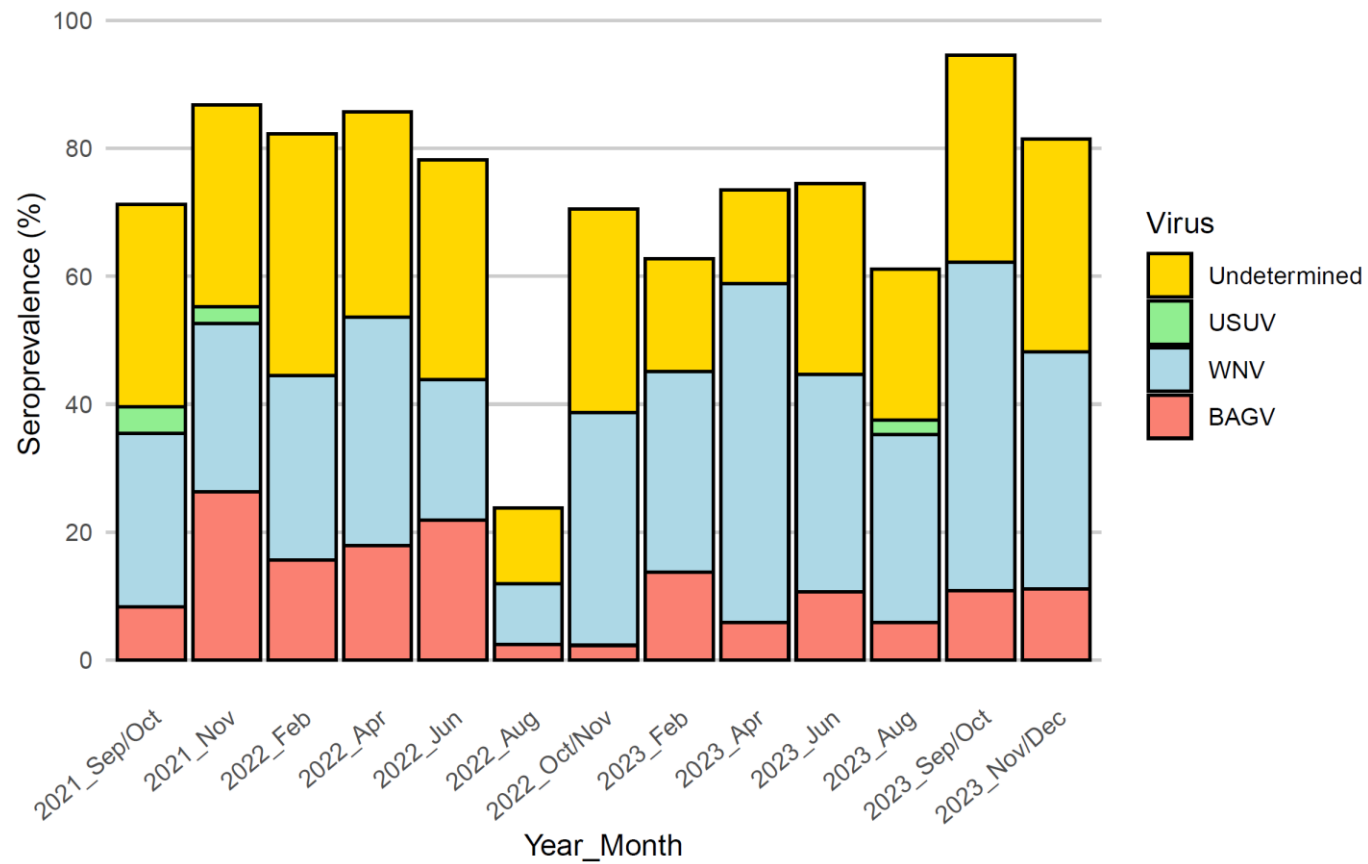

**Figure S1.** Specific antibody seroprevalences by micro virus neutralization (VNT) assay for *Alectoris rufa* sampled in Serpa. Blue for West Nile virus (WNV), green for Usutu virus (USUV), orange for Bagaza virus (BAGV) and yellow for unspecified antibodies (Undetermined).

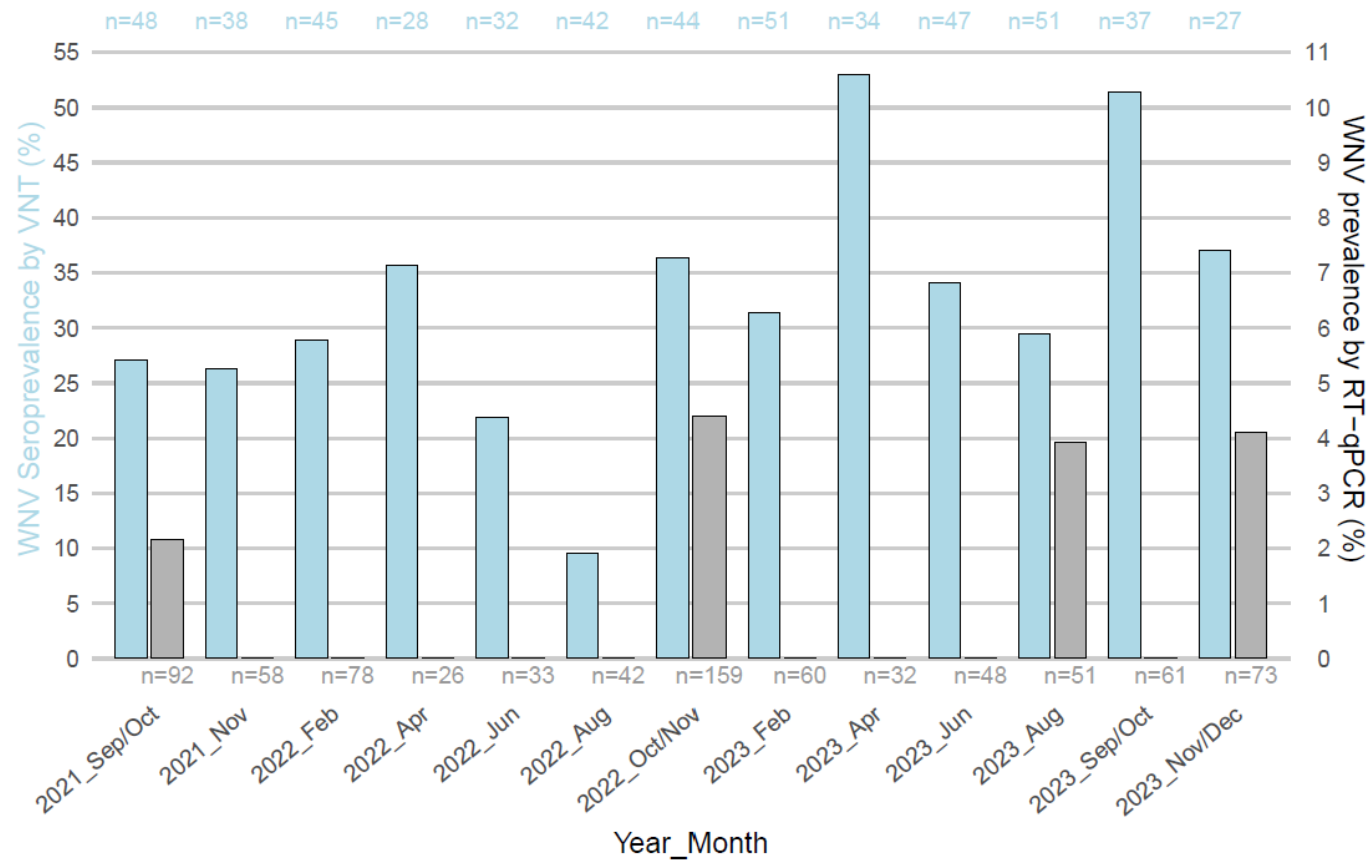

**Figure S2.** Seroprevalences (%) to West Nile Virus (WNV) by micro virus neutralisation (VNT) assay (scale on the left axis and blue coloured bars) and WNV prevalences (%) assessed by RT-qPCR (scale on the right axis and grey coloured bars) for *Alectoris rufa* sampled in Serpa. Total number of birds analysed by VNT is presented on top of the bars and total number of birds analysed by RT-qPCR is presented on the bottom.

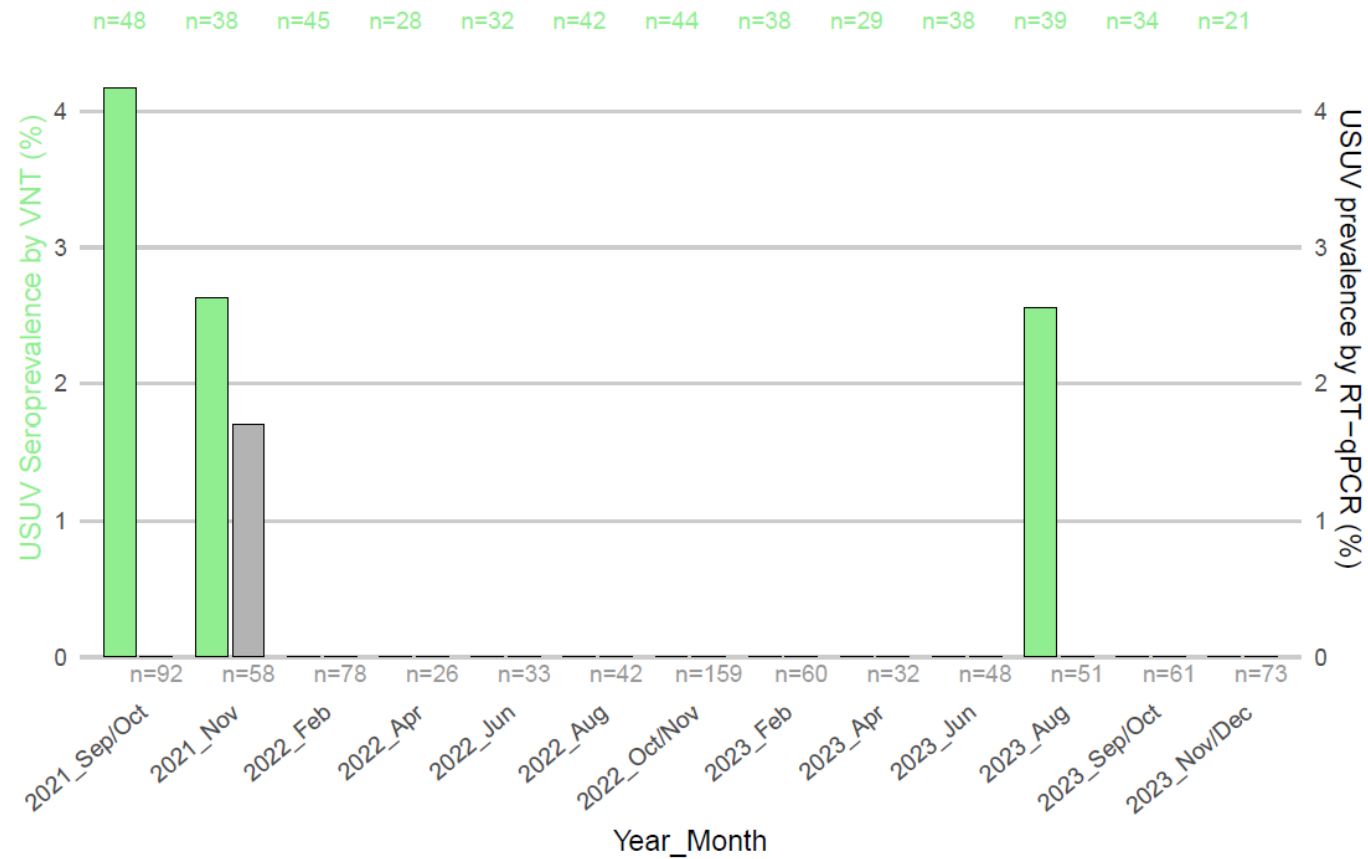

**Figure S3.** Seroprevalences (%) to Usutu virus (USUV) by micro virus neutralization (VNT) assay (scale on the left axis and green coloured bars) and USUV prevalences (%) assessed by RT-qPCR (scale on the right axis and grey coloured bars) for *Alectoris rufa* sampled in Serpa. Total number of individuals analysed by VNT is presented on top of the bars and total number of individuals analysed by RT-qPCR is presented on the bottom.

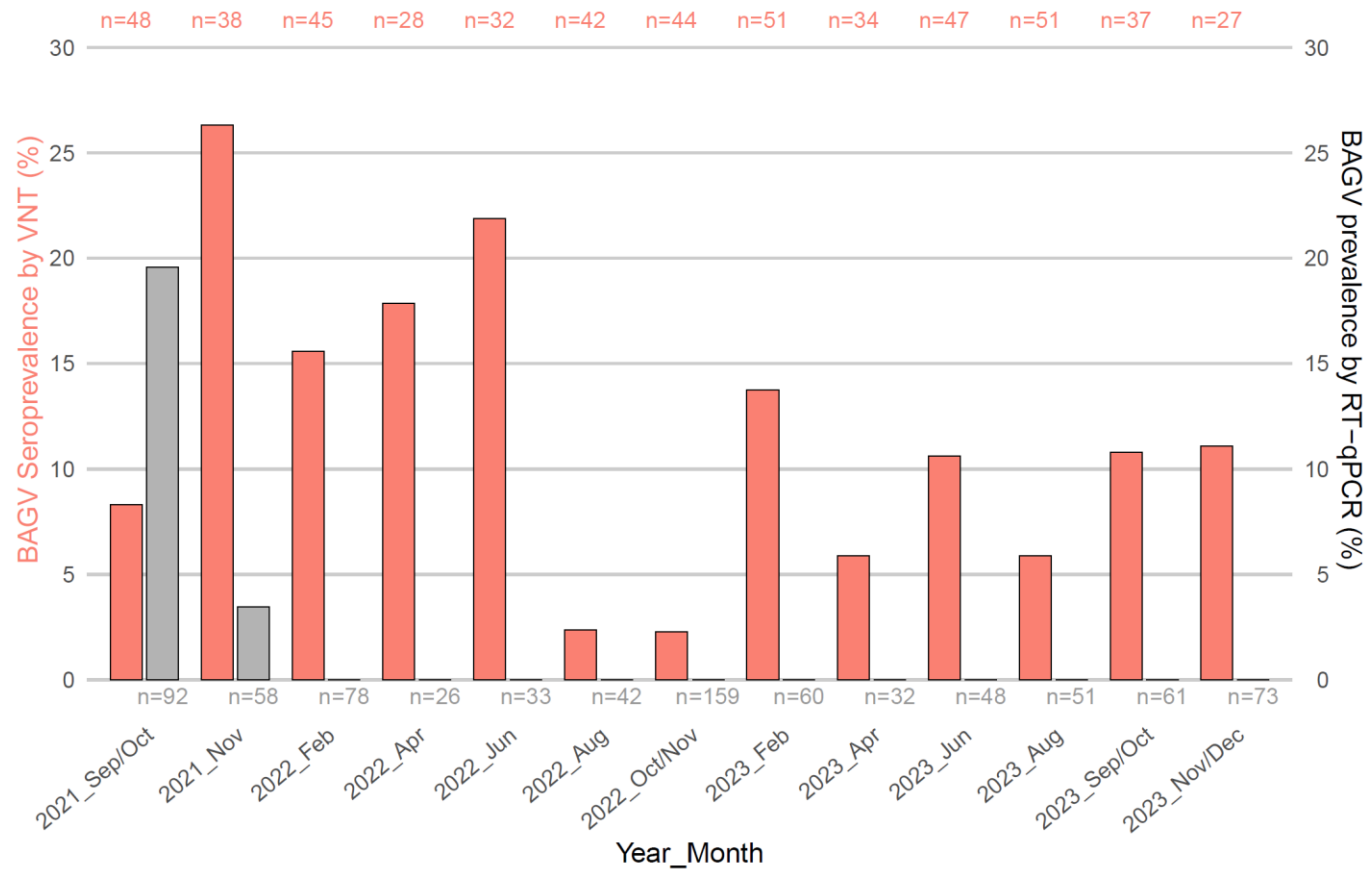

**Figure S4.** Seroprevalences (%) to Bagaza virus (BAGV) by micro virus neutralisation (VNT) assay (scale on the left and orange coloured bars) and BAGV prevalences (%) assessed by RT-qPCR (scale on the right and grey coloured bars) for *Alectoris rufa* sampled in Serpa. Total number of birds analysed by VNT is presented on top of the bars and total number of birds analysed by RT-qPCR is presented on the bottom.
